# Histone H1 protects telomeric repeats from H3K27me3 invasion in *Arabidopsis*

**DOI:** 10.1101/2020.11.28.402172

**Authors:** Gianluca Teano, Lorenzo Concia, Léa Wolff, Léopold Carron, Ivona Biocanin, Kateřina Adamusová, Miloslava Fojtová, Michael Bourge, Amira Kramdi, Vincent Colot, Ueli Grossniklaus, Chris Bowler, Célia Baroux, Alessandra Carbone, Aline V. Probst, Petra Procházková Schrumpfová, Jiří Fajkus, Simon Amiard, Stefan Grob, Clara Bourbousse, Fredy Barneche

**Affiliations:** Institut de biologie de l’Ecole normale supérieure (IBENS), Ecole normale supérieure, CNRS, INSERM, Université PSL, Paris, France; Sorbonne Université, CNRS, IBPS, UMR 7238, Laboratoire de Biologie Computationnelle et Quantitative (LCQB), 75005, Paris, France; Mendel Centre for Plant Genomics and Proteomics, Central European Institute of Technology, Masaryk University, Brno, Czech Republic; Laboratory of Functional Genomics and Proteomics, NCBR, Faculty of Science, Masaryk University, Brno, Czech Republic; Cytometry Facility, Imagerie-Gif, Université Paris-Saclay, CEA, CNRS, Institute for Integrative Biology of the Cell (I2BC), 91198, Gif-sur-Yvette, France; Department of Plant and Microbial Biology & Zürich-Basel Plant Science Center, University of Zürich, Switzerland; CNRS UMR6293, Université Clermont Auvergne, INSERM U1103, GReD, CRBC, Clermont-Ferrand, France; Université Paris-Saclay, 91190 Orsay, France

## Abstract

While the pivotal role of linker histone H1 in shaping nucleosome organization is well established, its functional interplays with chromatin factors along the epigenome are just starting to emerge. Here we first report that in *Arabidopsis*, as in mammals, H1 occupies Polycomb Repressive Complex 2 (PRC2) target genes where it favors chromatin condensation and H3K27me3 deposition. We further show that, contrasting with its conserved function in PRC2 activation at genes, H1 selectively prevents H3K27me3 accumulation at telomeres and large pericentromeric interstitial telomeric repeat (ITR) domains by restricting DNA accessibility to Telomere Repeat Binding (TRB) proteins, a group of H1-related Myb factors mediating PRC2 *cis* recruitment. This study unveils a mechanistic framework by which H1 avoids the formation of gigantic H3K27me3-rich domains at telomeric sequences and contributes to safeguard nucleus architecture.

Teano et al. report that that linker histone H1 and a group of H1-related telomeric proteins interplay to selectively influence the *Polycomb* repressive landscape at genes and telomeric repeats in *Arabidopsis*. These findings provide a mechanistic framework by which H1 influences the epigenome and nuclear organization in a sequence-specific manner.

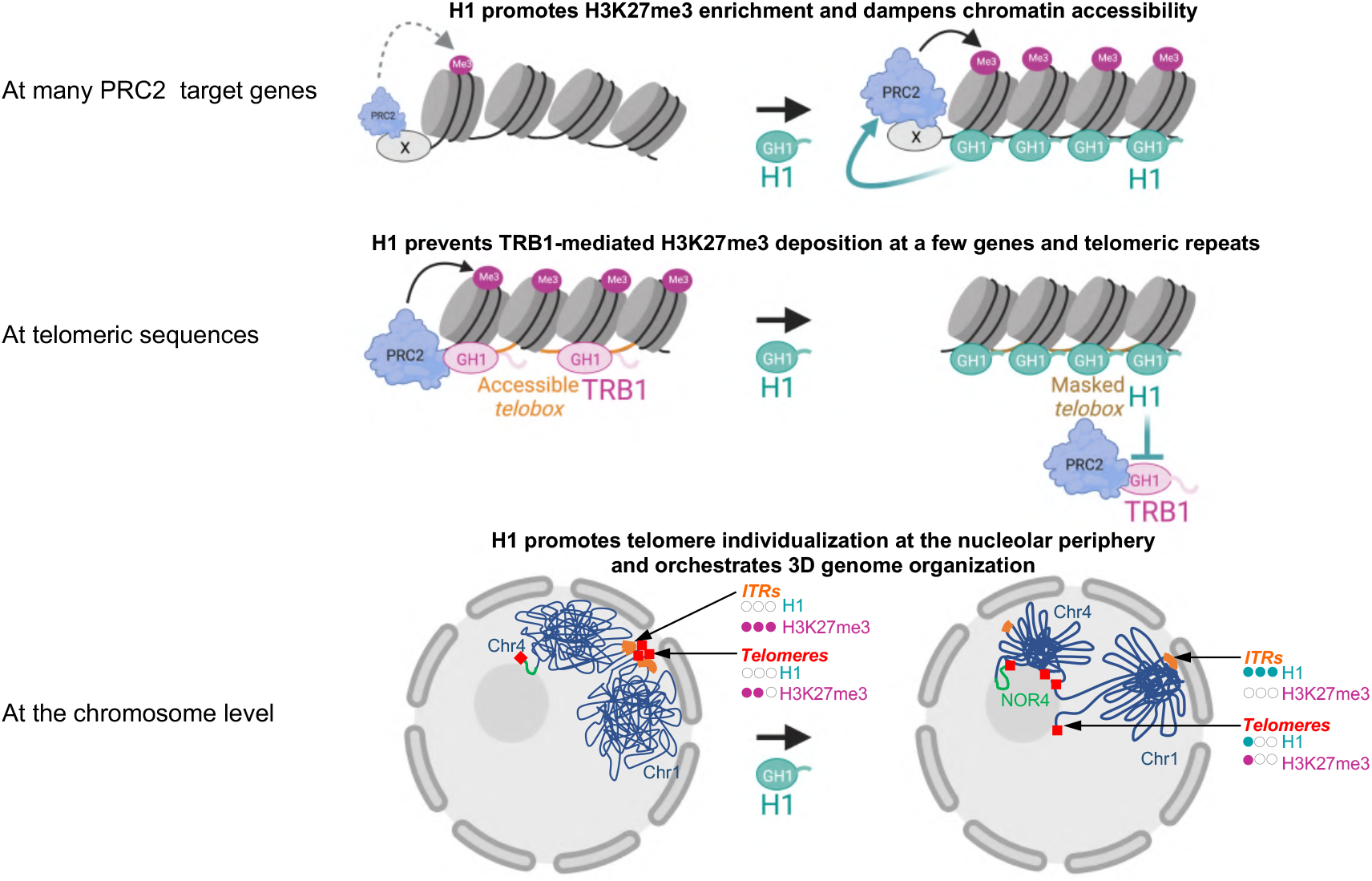

**Highlights:**

- H1 promotes PRC2 activity and limits accessibility at a majority of genes
- H1 prevents PRC2 activity at telomeric DNA sequences
- PRC2 repression is achieved by restricting accessibility to TRB proteins
- H1 orchestrates the spatial organization of telomeres and interstitial telomeres (ITRs)

## Introduction

Both local and higher-order chromatin architecture rely to a large extent on the regulation of nucleosome density and accessibility, in which linker histone H1 and *Polycomb* Repressive complexes 1 and 2 (PRC1/2) play distinct roles. H1 modulates nucleosome distribution by contacting the nucleosome dyad with its central globular (GH1) domain and binding linker DNA at the nucleosome entry and exit sites with its disordered carboxy-terminal domain. This indirectly contributes to dampen transcriptional activity by affecting the accessibility of transcription factors and RNA polymerases to chromatin but also through interactions with histone and DNA modifiers (reviewed in ^1–3^).

*Polycomb* Group activity is another determinant of chromatin organization that extensively regulates transcriptional activity, cell identity, and differentiation in metazoans ^4,5^, plants ^6^ and unicellular eukaryotes ^7^. While H1 incorporation directly influences the physicochemical properties of the chromatin fiber, *Polycomb* Repressive Complex 1 (PRC1) and 2 (PRC2) display enzymatic activities mediating histone H2A Lysine monoubiquitination (H2Aub) and histone H3 Lysine 27 trimethylation (H3K27me3), respectively ^4,5^. In metazoans, chromatin of PRC target genes is highly compacted ^8–10^, a feature thought to hinder transcription (reviewed in ^5,11^). PRC2 can favor chromatin compaction either by promoting PRC1 recruitment or through its subunit Enhancer of Zeste Homolog 1 (Ezh1) in a mechanism not necessarily relying on the H3K27me3 mark itself ^12^.

Mutual interplays between H1 and PRC2 activity first emerged *in vitro*. Human H1.2 preferentially binds to H3K27me3-containing nucleosomes ^13^ while, *vice versa*, human and mouse PRC2 complexes display substrate preferences for H1-enriched chromatin fragments. The latter activity is stimulated more on di-nucleosomes than on mono- or dispersed nucleosomes ^14,15^. *In vivo*, recent studies unveiled that H1 is a critical regulator of H3K27me3 enrichment over hundreds of PRC2 target genes in mouse cells ^16,17^. Chromosome conformation capture (Hi-C) analysis of hematopoietic cells ^16^, germinal centre B cells ^17^, and embryonic stem cells ^18^ showed that H1 triggers distinct genome folding during differentiation in mammals. These major advances raise the question of the mechanisms enabling H1 sequence-specific interplays with PRC2 activity in chromatin regulation and their evolution in distinct eukaryotes.

In *Arabidopsis thaliana*, two canonical linker histone variants, H1.1 and H1.2, represent the full H1 complement in most somatic cells ^19–21^. These two linker histones, hereafter referred to as H1, are enriched over heterochromatic transposable elements (TEs) displaying high nucleosome occupancy, CG, CHG and CHH methylation as well as H3K9 dimethylation ^22,23^. While also contributing to CG methylation mediated gene silencing ^24^, H1 is less abundant over expressed genes ^22,23^. As in mammals, *Arabidopsis* H1 incorporation is thought to dampen RNA Pol II transcription, an effect that also applies in plants to RNA Polymerase Pol IV that produces short interfering RNAs (siRNAs) ^25^. *Arabidopsis* H1 also restricts accessibility to DNA methyltransferases and demethylases that mediates gene or TE silencing ^26,27,27–30^. This process is counter-balanced by incorporation of the H2A.W histone variant presumably competing with H1 for DNA binding through its extended C-terminal tail ^31^. Interestingly, recent studies suggested that H1 dynamics may impact PRC2 activity during *Arabidopsis* development. The first piece of evidence comes from the observation that H1 is largely absent from the vegetative cell nucleus of pollen grain and is degraded during spore mother cell (SMC) differentiation at the onset of heterochromatin loosening and H3K27me3 reduction ^26,32–34^. The second evidence comes from the observation that *H1* loss-of-function mutant nuclei display a ^~^2-fold lower H3K27me3 chromatin abundance, while a few discrete H3K27me3 subnuclear foci of undetermined nature displayed increased H3K27me3 signals ^35^. Hence, despite evidence that variations in H1 abundance mediate epigenome reprogramming during plant development, there is no information on how H1 interplays with PRC2 activity and on the consequences of this interaction on the chromatin landscape and topology in these organisms.

Here, we profiled H3K27me3 in *h1* mutant plants and found that, whilst a majority of genes expectedly lost H3K27me3, telomeres and pericentromeric interstitial telomeric regions (ITR/ITS) were massively enriched in this mark. We identified that H1 prevents PRC2 activity at these loci by hindering the binding of Telomere Repeat Binding (TRB) proteins, a group of H1-related proteins with extra-telomeric function in PRC2 recruitment ^36,37^. H1 safeguards telomeres and ITRs against excessive H3K27me3 deposition and preserves their topological organization. Collectively, our findings led us to propose a mechanism by which H1 orchestrates *Arabidopsis* chromosomal organization and contributes to the control of H3K27me3 homeostasis between structurally distinct genome domains.

## Results

### H1 is abundant at H3K27me3-marked genes and reduces their chromatin accessibility

To assess the relationships between H1, PRC2 activity and chromatin accessibility, we first compared the genomic distribution of H3K27me3 with that of H1.2, the most abundant canonical H1 variant in *Arabidopsis* seedlings ^22^. To maximize specificity, we used an GFP-tagged version of H1.2 expressed under the control of its endogenous promoter ^22^. In agreement with previous studies in several eukaryotes ^22,23,38,39^, this showed that H1.2 covers most of the *Arabidopsis* genome without displaying clear peaks. Yet, a closer examination revealed that, as compared to genes and to TEs that are not enriched in H3K27me3 ^40–42^, H1 level was higher at coding genes marked by H3K27me3, especially towards their 5’ region (Figure 1A, S1A, S2A).

**Figure 1.**
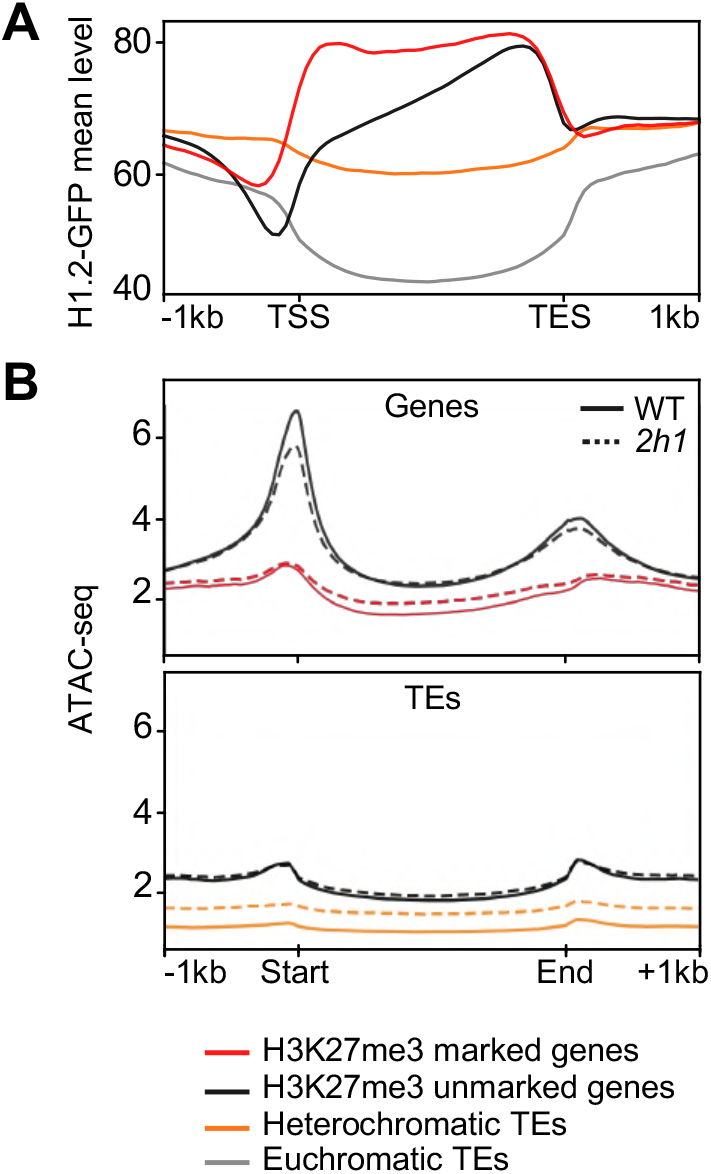
H1.2-GFP is enriched at PRC2-target genes where it contributes to restrain DNA accessibility. **A.** H1.2-GFP mean read coverage at protein-coding genes and TEs. **B.** ATAC-seq analysis of chromatin accessibility of genes and TEs described in (A) in WT (plain lines) and *2h1* (dashed lines) nuclei. Chromatin accessibility is estimated as read coverage. TSS, Transcription Start Site. TES, Transcription End Site. In (A-B), H3K27me3-marked genes (n=7542) are compared to all other annotated protein-coding genes. Heterochromatic versus euchromatic TEs were defined previously ^43^. The plots represent the mean of three (H1.2-GFP) or two (ATAC-seq) independent biological replicates.

Having found that H1 is enriched at PRC2 marked genes, we tested whether it contributes to regulate chromatin accessibility using Assay for Transposase-Accessible Chromatin followed by sequencing (ATAC-seq) in nuclei of WT and *h1.1h1.2* double mutant plants (hereby named *2h1* for short). As previously reported in WT plants ^44^, H3K27me3-marked genes displayed low chromatin accessibility as compared to non-marked genes, which are usually expressed and typically display a sharp ATAC peak at their TSS corresponding to the nucleosome free region (Figure 1B). In *2h1* nuclei, gene body regions of H3K27me3-marked loci displayed a significant increase in accessibility (Figure 1B and S2B-C). Hence, H1 tends to abundantly occupy PRC2 target gene bodies where it has a minor but detectable contribution in restricting chromatin accessibility.

### H1 promotes H3K27me3 enrichment at a majority of PRC2 target genes whilst protecting a few genes displaying specific sequence signatures

To determine at which loci H1 influences PRC2 activity, we profiled the H3K27me3 landscape in WT and *2h1* seedlings. To enable absolute quantifications despite the general reduction of H3K27me3 in the mutant nuclei, we employed ChIP-seq with reference exogenous genome (ChIP-Rx) by spiking-in equal amounts of Drosophila chromatin to each sample ^45^ (Additional File 1). Among the ^~^7,500 genes significantly marked by H3K27me3 in WT plants (Figure S3D), more than 4,300 were hypomethylated in *2h1* plants (Figure 2A-C, Figure S3A-C and Additional File 1). Hence, general loss of H3K27me3 in *2h1* seedlings identified by immunoblotting and cytology ^35^ results from a general effect at a majority of PRC2 regulated genes. It is noteworthy that ^~^85% of the genes marked by H3K27me3 in WT plants were still significantly marked in *2h1* plants (Figure S3D). Hence, H1 is required for efficient H3K27me3 maintenance or spreading but less for PRC2 seeding. Our RNA-seq analysis showed that genes encoding PRC1/PRC2 subunits are not downregulated in *2h1* plants, excluding indirect effects resulting from less abundant PRC2 (Additional File 2). Unexpectedly, we also found that ^~^500 genes were hyper-marked or displayed *de novo* marking in *2h1* plants (Figure 2A-C and Additional File 1).

**Figure 2.**
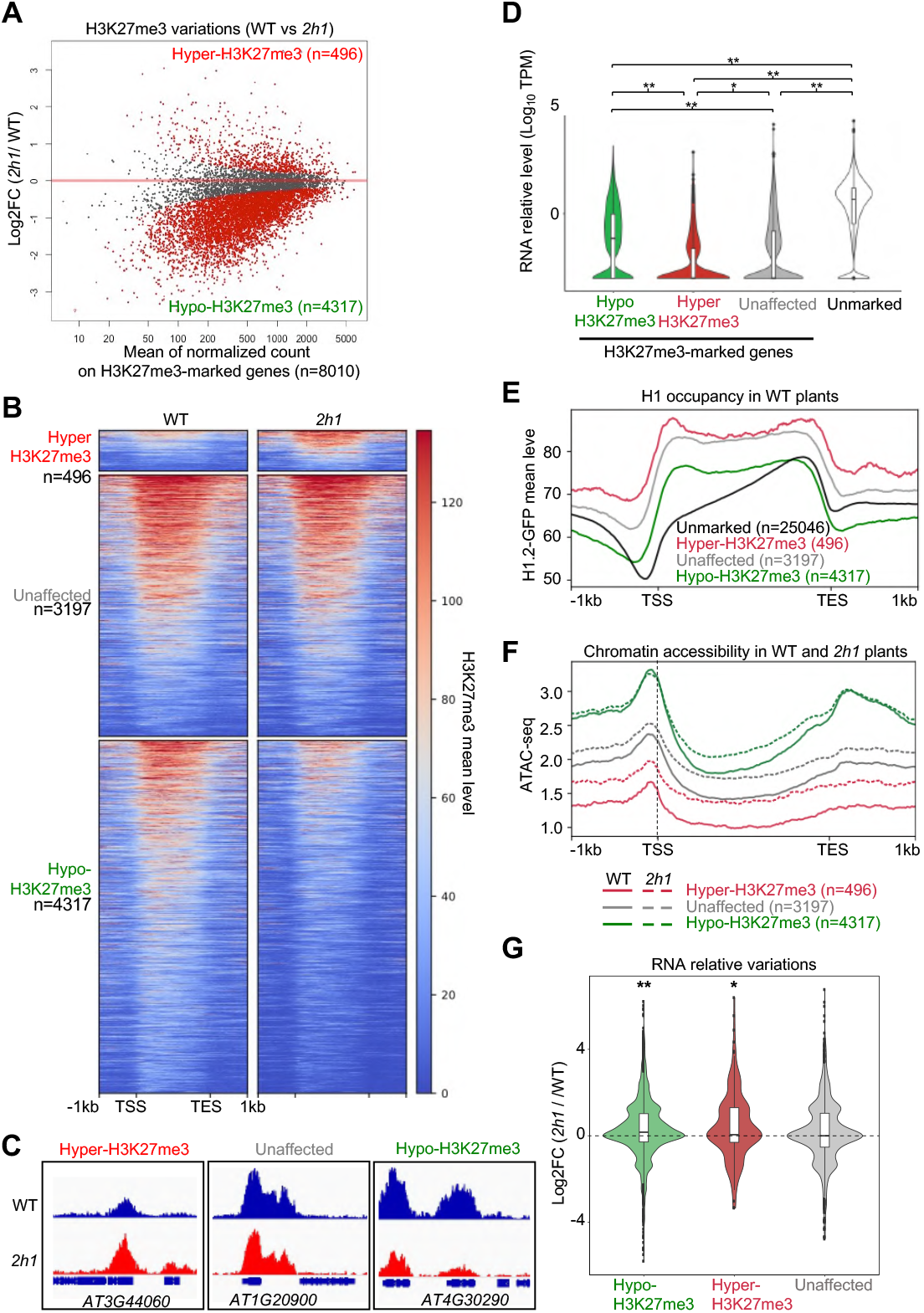
H1 influences H3K27me3 marking, chromatin accessibility and expression of PRC2-target genes. **A.** Identification of differentially marked genes using spike-in normalized DESeq2 ChIP-seq analysis identifies low H3K27me3 levels over a majority of the PRC2 target genes in *2h1* plants. All genes displaying an H3K27me3-enriched domain in WT or *2h1* plants (according to MACS2 peak detection, see Methods) are individually shown as dots. Red dots, differentially marked genes (FDR < 0.01). **B.** H3K27me3 profiles along all genes significantly marked in WT or *2h1* plants. Genes are grouped according to differential analysis in (A) and ranked within each group according to mean H3K27me3 levels. **C.** H3K27me3 profile of representative genes of the three sets identified in (A) exemplifying the general tendency of PRC2-target genes to keep a weak H3K27me3 domain in *2h1* plants. **D.** Transcript levels in WT seedlings. The values represent RNA-seq Log10 TPM values. The embedded box plots display the median while lower and upper hinges correspond to the first and third quartiles. * and ** indicate p-values below 10^-9^ and 10^-15^, respectively, according to a Wilcoxon rank test. **E.** H1.2-GFP ChIP-seq profiling on the indicated gene sets (mean read coverage). **F.** ATAC-seq analysis of the indicated gene sets. ATAC-seq data are presented as in Figure 1B using mean read coverage. **G.** Transcript level variations between WT and *2h1* plants in the same three gene sets. The values represent mRNA Log_2_ fold changes. The embedded box plots display the median while lower and upper hinges correspond to the first and third quartiles. * and ** indicate p-values below 5%, 1%, respectively, according to a Student t-test. ChIP-Rx, ATAC-seq and RNA-seq data correspond to two biological replicates, each, and H1.2-GFP ChIP-seq to three biological replicates.

To determine whether the hypo/hyper/unaffected gene sets had different functional properties, we inspected their transcript level. Hyper-marked genes correspond to the least expressed gene category in WT plants whereas many hypo-marked genes are significantly more expressed than unaffected genes (Figure 2D). Functional categorization of hypo-marked genes notably identified an over-representation of genes involved in transcriptional regulation and meristem maintenance (Figure S5A). These classifications are consistent with former reports of PRC2 repressing these biological processes ^46^. In contrast, a feature of the hyper-marked gene set is the presence of TE or TE-gene annotations (Figure 2F, S5, Additional File 1). Hence, we concluded that H1-mediated PRC2 activation ^14,15,16^ is conserved in plants, but in *Arabidopsis* this property is contrasted by a heretofore-unsuspected negative effect at a minority of poorly expressed genes sometimes displaying TE features.

*In vitro*, PRC2 activity was proposed to be favored by local H1 abundance and/or at densely organized nucleosome arrays ^14^. Instead, we found that hypo-marked genes tend to display lower H1 level, lower nucleosome occupancy, and to be more accessible and expressed than other genes marked by H3K27me3 (Figure 2E-G and S4). We therefore tested nucleosome density using ChIP-seq profiling of histone H3, confirming that hypo-marked genes display lower nucleosome occupancy than other marked gene categories (Figure S4C). Collectively, analysis of the hypo-marked loci suggests that chromatin of the corresponding genes is not sufficiently nucleosome dense to favor PRC2 *cis* activity when H1 is absent.

We further explored whether the specific influence of H1 on H3K27me3 enrichment at genes could rely on a sequence-dependent mechanism, especially at hyper-marked genes since they do not incur the conserved H1-mediated PRC2 activation. In contrast to the promoter sequences of the hypo-marked genes in which no such motif is significantly over-represented, we identified three enriched motifs in the hyper-marked gene set (Figure S5C). A poly-A motif is present in 84% of the gene promoters, and the AAACCCTA telomeric motif, referred to as *telobox* ^47,48^ that serves as *Polycomb Response Elements* (*PREs*) in plants ^36,37^ is found in 17% of them (Additional File 1). Based on these observations, we conclude that the capacity of H1 to counteract H3K27me3 enrichment at a small gene set presumably involves specific sequence features.

### H1 contributes to define accessibility and expression of PRC2 target genes

To get insights into the functional consequences of H1 loss at genes where it either promotes or dampens H3K27me3 enrichment, we compared the chromatin accessibility and transcript levels of these gene sets in WT and *2h1* nuclei. ATAC-seq profiling showed that hypo-marked gene bodies were significantly more accessible in the mutant line (Figure 2F and S4B), thereby correlating with reduced H3K27me3 levels. Despite H3K27me3 gain accessibility of hyper-marked genes was increased in *2h1* plants, but it remained very low (Figure S4B). Conservation of this function in both gene categories indicates that H1 incorporation reduces chromatin accessibility of *Arabidopsis* PRC2-target genes independently of its influence on H3K27me3 enrichment.

Confirming previous reports ^23,35^, our RNA-seq analysis showed that *H1* loss-of-function triggers minor gene expression changes (Additional File 2). Yet, we identified a significant tendency for increased transcript levels of the H3K27me3 hypo- and hyper-marked genes set in the *2h1* line (Figure 2G). Taken together, these analyses showed that, at a majority of PRC2 target genes, H1 depletion triggers H3K27me3 loss associated to a moderate increase in DNA accessibility and expression.

### H1 prevents H3K27me3 invasion over a specific family of heterochromatic repeats

Considering the observed H3K27me3 enrichment at a few TE-related genes in *2h1* plants, we extended our analysis to TEs, which typically lack H3K27me3 in *Arabidopsis*^49,50^. This revealed that 1066 TEs are newly marked by H3K27me3 in *2h1* plants, most frequently over their entire length, thereby excluding *a priori* the possibility that H3K27me3 TE enrichment is due to spreading from neighboring genes (Figure 3A). We clustered H3K27me3-marked TEs into two groups, *TE cluster 1* and *TE cluster 2* displaying high and low H3K27me3 enrichment, respectively (Figure 3A). While *TE cluster 2* (n=850) is composed of a large variety of TE families, *TE cluster 1* (n=216) mostly consists of *ATREP18* (189 elements) annotated in the TAIR10 genome as *Unassigned* (Figure 3B). In total, *TE cluster 1* and *2* comprise 60% of all *Arabidopsis ATREP18* elements including many of the longest units (Figure S6A). A second distinguishing feature of *TE cluster 1* elements is their elevated H1 and H3 occupancy (Figure 3C, S6B and S7A). Accordingly, *TE cluster 1* and more generally *ATREP18* elements are strongly heterochromatic with elevated H3K9me2 nucleosome occupancy, cytosine methylation and very low chromatin accessibility (Figure 3D, S6C-D, Figure S7B,G). Taken together, these observations indicate that H1 prevents H3K27me3 accumulation over a set of H1-rich, heterochromatic, and highly compacted repeats, which contrasts with its positive influence on H3K27me3 marking over thousands of PRC2-target genes.

**Figure 3.**
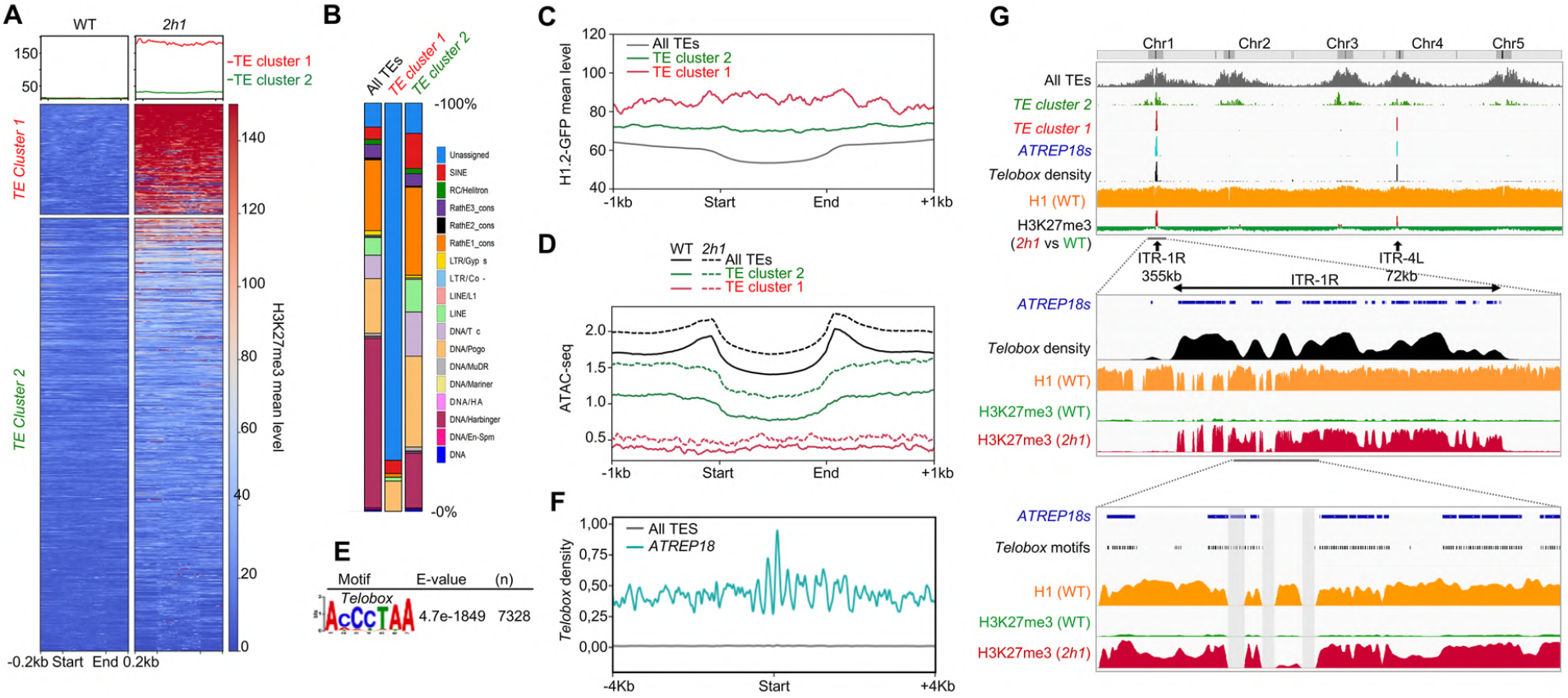
H1 hinders H3K27me3 enrichment at two pericentromeric ITR blocks spanning more than 420 kb. **A.** Hyper-marked TEs were clustered into two groups according to H3K27me3 levels after spike-in normalization defining two TE clusters of 216 and 850 TEs, respectively. H3K27me3 profiles over all *TE Cluster 1* and *2* elements are ranked in each group according to H3K27me3 mean spike-in normalized coverage. **B.** Relative TE superfamily composition of H3K27me3-enriched TEs. *TE cluster 1* comprises a strong over-representation of “Unassigned” annotations mainly corresponding to *ATREP18* elements, while *TE cluster 2* elements correspond to a wide variety of TE super-families. **C.** *TE Cluster 1* elements display high H1 occupancy. The plot represents H1.2-GFP mean read coverage over the indicated repertoire of TEs and repeats. **D.** Chromatin accessibility of *TE cluster 1* elements remains very low in *2h1* nuclei. ATAC-seq data are presented as in Figure 1B using mean read coverage. **E.** Motif enrichment search identified an over-representation of *telobox* motifs in *TE cluster 1* sequences. E-values were calculated against all TE sequences. **F.** *ATREP18* repeats display outstanding density and a distinct pattern of *telobox* motifs as compared to whole set of annotated TEs. The plot represents the density of perfect *telobox* sequence motifs in all *ATREP18s* as compared to all TEs within 50bp bins. **G.** Chromosome distribution of H3K27me3 defects in *2h1* plants and their link to *ATREP18*, *TE Cluster 1* and *TE Cluster 2* elements. The sharp peaks of *telobox* density in the pericentromeres of chromosome 1 and 4 correspond to Interstitial Telomeric Repeat ITR-1R and ITR-4L. Chromosome 1 pericentromeric region displays a sharp overlap between *2h1*-specific H3K27me3 enrichment and the *telobox*-rich ITR-1R. Bottom panel, shaded boxes correspond to blacklisted TAIR10 genome sequences (see Methods). Complementary profiles over ITR-4L and other interspersed elements from *TE Cluster 2* are shown in Figure S8.

Noteworthy, while MNase-seq analyses^22,23^ and our ATAC-seq data showed that heterochromatic TEs tend to be more accessible in *2h1* nuclei, the chromatin of *TE cluster 1* and *ATREP18* repeats remained very poorly accessible despite H1 loss (Figure 3D and S6C,D). Hence, chromatin “inaccessibility” of *TE Cluster 1* elements is either H1-independent or compensated by other mechanisms, possibly the increased PRC2 local activity.

### Repeats gaining H3K27me3 in *2h1* plants are parts of two large blocks of pericentromeric telomeric repeats

Aiming at determining the features potentially leading to a selective role of H1 at *TE Cluster 1* elements, we first envisaged that H1 could locally prevent conversion of the H3K27me1 heterochromatic mark into H3K27me3. However, analysis of public datasets ^51^ showed that, as compared to other TEs, H3K27me1 is not particularly abundant at *TE cluster 1* or at *ATREP18* elements, therefore ruling out this first hypothesis (Figure S7C). We then explored the possibility that H1 could rather favor H3K27me3 de-methylation. Examination of the H3K27me3 profile in loss-of-function plants for the three major histone H3K27me3 demethylases EARLY FLOWERING 6 (ELF6), RELATIVE OF ELF 6 (REF6), and JUMONJI 13 (JMJ13) ^52^ showed no H3K27me3 increase at *TE cluster 1* elements (Figure S7E) nor at hyper-marked genes (Figure S7F). This led us to rule out the hypothesis that in WT plants H3K27me3 could be regulated at these loci though active erasure. Last, considering the tendency for cytosine methylation to be mutually exclusive with H3K27me3 deposition in *Arabidopsis* ^53–55^, we envisioned that H3K27me3 enrichment at *TE Cluster 1* may indirectly result from decreased DNA methylation induced by H1 loss. Examination of cytosine methylation patterns of *TE Cluster 1* elements in *2h1* plants oppositely showed an increase in CG, CHG and CHH methylation (Figure S7G). We did not ascertain whether methylated cytosines and H3K27me3-containing nucleosomes co-occur at individual *TE Cluster 1* chromatin fragments, yet this observation ruled out that H1 indirectly hinders PRC2 activity at these loci by promoting cytosine methylation, a possibility that would have been supported if an opposite effect was observed.

Having not found evidence for indirect roles of H1 on H3K27me3 marking at *TE Cluster 1*, we concluded that H1 hinders PRC2 recruitment or activity at these repeats, and this despite their densely-packed chromatin organization theoretically constituting an excellent substrate. As previously done for hyper-marked genes, we therefore tested whether *TE Cluster 1* elements are distinguishable from other TEs by specific DNA motifs. MEME search identified a prominent sequence signature, the *telobox* motif, which we had already identified in 17% of the hyper-marked genes (Figure 3E, S5C). As compared to all other TEs, *teloboxes* were found to be ^~^100-fold more densely represented in *ATREP18* elements as compared to all TEs (Figure 3E-F). With 7328 *telobox* motifs, *TE cluster 1* contains ^~^53% of the whole TAIR10 *telobox* repertoire (Additional File 1). Hence, if not considering proper telomeres that span 2 to 5 kb at the end of each chromosome ^56,57^, *TE Cluster 1* repeats display the majority of telomeric motifs of *Arabidopsis* genome and the strongest propensity to attract PRC2 activity upon H1 loss.

Remarkably, these two properties can be seen at a chromosome scale by contrasting the genome distribution of *telobox* density and of H3K27me3 differential marking, since about 95 % of *TE Cluster 1* elements cluster within two outstandingly *telobox*-rich regions situated in the pericentromeres of chromosomes 1 and 4 (Figure 3G and S8,S9). Given this characteristic, we consider these domains as two of the nine *Arabidopsis* genome loci proposed to constitute ITRs ^58,59^, hereby referred to as ITR-1R and ITR-4L of ^~^355 kb and ^~^72 kb, respectively. In agreement with the description of ITRs in plants and vertebrates ^60,61^, *ATREP18* elements that constitute most of these domains display a high density in *telobox* motifs frequently organized as small clusters (Figure 3G and Figure S9). Further supporting their telomeric evolutionary origin, *ATREP18s* encode no open reading frame or other TE features, are mostly oriented on the same DNA strand, and tandemly organized (nearly 90 % of them being positioned within 1 kb of each other, Figure S10), hence they do not constitute *stricto sensu* TEs. Ectopic H3K27me3 deposition was also found at several interspersed elements of *TE cluster 2* located in all pericentromeric regions outside these two ITR blocks (Figure S8B), but our main conclusion is that H1 abundantly occupies two large blocks of pericentromeric ITRs where it prevents H3K27me3 marking.

### H1 influences telomere chromatin composition and sub-nuclear positioning

Considering that telomeres display hundreds perfect *telobox* motifs, the question arose whether, similarly to ITRs, H1 also prevents H3K27me3 deposition at chromosome ends. Because the perfect continuum of terminal telomeric motifs is not suited for quantitative NGS analyses, ChIPs were analyzed through hybridization with radioactively labeled concatenated telomeric probes ^62^. H3K27me3 ChIP dot-blots led to the estimation that telomeres display an average ^~^4-fold more H3K27me3 enrichment in *2h1* as compared to WT plants, independently of detectable changes in nucleosome occupancy probed by anti-H3 ChIP dot-blot (Figure 4A and S11A).

**Figure 4.**
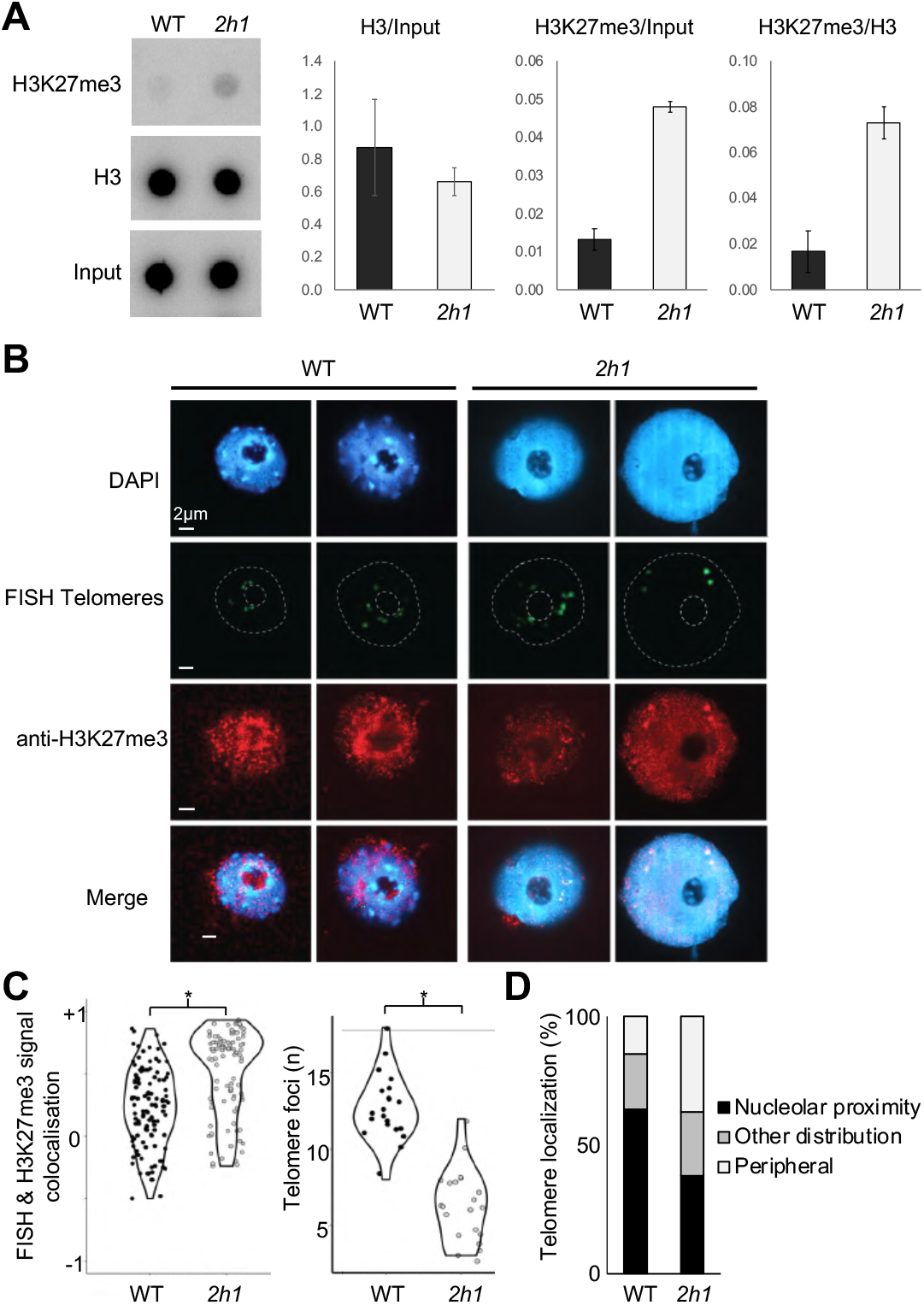
H1 influences both H3K27me3 enrichment and sub-nuclear organization of telomeres. **A.** Increased H3K27me3 level at telomeres in *2h1* plants. H3 ChIP signal is used as a proxy of nucleosome occupancy. ChIPs were followed by dot-blot hybridization with a labeled telomeric probe. Data are the mean of two biologically and technically replicated experiments +/-SE. A second biological replicate is shown in Figure S11. **B.** Most telomeric loci are enriched in H3K27me3 and re-distributed toward the nucleus periphery in *2h1* plants. Representative collapsed Z-stack projections of cotyledon nuclei subjected to H3K27me3 immunolabeling and telomere DNA FISH are shown. Blue, DAPI DNA counterstaining; Green, telomere FISH signals; Red, H3K27me3 immunolabeling. **C.** Quantification of sub-nuclear telomeric signal properties. * indicates p-Values <1.6e–07 Wilcoxon signed-rank test. **D.** Quantification of nuclei classes displaying different patterns in telomere sub-nuclear localization. Number and position of telomeric foci were determined in two independent biological replicates (n>20 each).

To assess whether H3K27me3 enrichment concerns a few telomeres or affects them all, we explored its occurrence in intact nuclei using H3K27me3 immunolabeling combined with telomere Fluorescence In Situ Hybridization (DNA FISH). Consistent with our ChIP-blot analysis, most telomeric foci were enriched with H3K27me3 in *2h1* nuclei, with 2-to-4 telomere foci frequently presenting outstandingly strong H3K27me3 signals (Figure 4B-C). We could not ascertain whether some of these strong signals correspond to cross-hybridizing pericentromeric ITRs, but their frequent positioning near to the nuclear periphery may point out to the latter hypothesis. Indeed, in *2h1* nuclei telomeric foci were frequently re-distributed toward the nucleus periphery, thereby contrasting with the ‘telomere rosette model’ proposed by Fransz and co-worker (2002) ^63^ first establishing that telomeres cluster around the nucleolus (Figure 4D). In addition, the number of telomere foci was reduced in the mutant nuclei (Figure 4B-C), indicating that H1 not only prevents accumulation of H3K27me3 at ITRs and at most telomeres but is also required for the sub-nuclear organization and proper individualization of these domains.

### H1 promotes heterochromatin packing but attenuates ITR insulation and telomere-telomere contact frequency

To better understand the altered telomere cytogenetic patterns of *2h1* nuclei and to extend our analysis to ITR topology, we employed *in situ* chromosome conformation capture (Hi-C) of dissected cotyledons, composed of 80% mesophyll cells, which enabled to reach high resolution (Figure S12, S13A).

In agreement with previous reports ^64–69^ WT plants displayed frequent interactions within and between pericentromeric regions, which reflect packing of these domains within so-called ‘chromocenter’ conspicuous structures (Figure 5A, 5B, S13A, S14A). Loosening of these heterochromatic domains in *2h1* mutant nuclei, formerly observed by microscopy ^23,35^, was expectedly identified as a more steep decay with distance ^70^ and lower long-range interaction frequency within pericentromeric regions (Figure 5A, 5B, S14). Yet, this tendency appears to be a general trend in the mutant nuclei since it was also observed for chromosome arms. As also seen in *crwn* and *condensin* mutants^71^, in a matrix of differential interaction frequency between WT and *2h1* nuclei these prominent defects are also visible as blue squares surrounding the centromeres, mirrored by increased interaction frequency between pericentromeric regions and their respective chromosome arms (i.e., red crosses along chromosome arms) (Fig. 5C).

**Figure 5.**
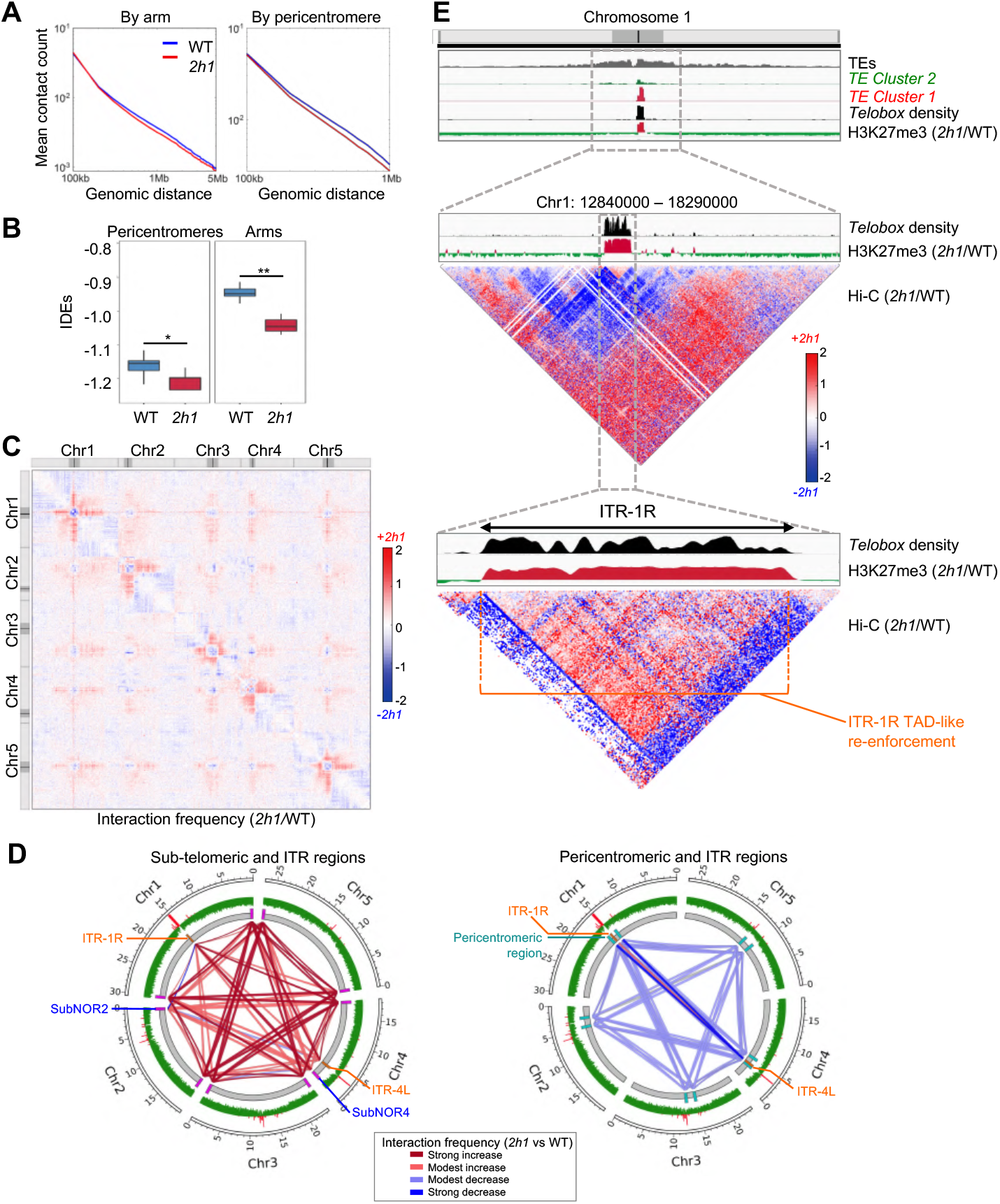
H3K27me3 accumulation at ITRs and at telomeres associates with ITR insulation and more frequent telomere-telomere interactions. **A.** Mean contact count as a function of genomic distance for all chromosome arms at a 100 kb resolution. **B.** Distribution of Interaction Decay Exponents (IDEs) determined at a 100 kb resolution for chromosome arms and pericentromeric regions of WT and *2h1* nuclei. Median IDE values of chromosome arms and pericentromeres were determined as −0.95/-1.16 in WT and −1.05/-1.2 in *2h1* nuclei, respectively. * and ** indicated pairwise Wilcoxon rank-sum test comparison P-values of 0.076 and 0.001, respectively. **C.** Relative difference of interaction frequency between WT and *2h1* plants. The Log_2_ values of observed/expected (O/E) interaction frequency of the five chromosomes in *2h1* versus WT are shown at a 100 kb resolution. Regions in red have more frequent contacts in *2h1* than in WT plants while regions in blue have less. Pericentromeric regions are depicted in dark grey on the schematic chromosomes 1. **D.** H1 reduces the frequency of long-distance interactions between chromosome ends. Circos-plots depict variations in inter-chromosomal interaction frequencies between telomere-proximal, pericentromeric, ITR-1R and ITR-4L 100 kb domains. Yellow boxes, ITR regions. External green/red track, H3K27me3 variations in *2h1* versus WT plants (Log2 ratio). Magenta boxes, telomere proximal regions and SubNOR2 or SubNOR4. **E.** Reduced frequency of intra-pericentromeric O/E interactions in *2h1* mutant nuclei is contrasted by TAD re-enforcement of the H3K27me3-enriched ITR-1R 355 kb block. Top panel, location of ITR-1R in chromosome 1. Middle panel, magnification of the region surrounding chromosome 1 pericentromeres at a 10 kb resolution. Bottom panel, magnification of the pericentromere-imbedded ITR-1R at a 2 kb resolution. Strong and modest increase correspond to Log2FC>1 and Log2FC 0.35/1, respectively; modest and strong decrease correspond to Log2FC −0.33/-0.65 and Log2FC<-0.65, respectively. Quantitative analyses are shown in the complementary Figure S14. All Hi-C analyses combine three independent biological replicates.

Having identified large-scale defects of chromosome organization in *2h1* mutant nuclei, we then focused on telomere-telomere interaction frequency. Because telomeres are not included in the TAIR10 reference genome, we used the most sub-telomeric 100 kb sequences of each chromosome end as a proxy to estimate telomere long-distance interactions, and these were controlled using an internal 100-kb region of each pericentromeric region as well as 100-kb regions randomly chosen in distal chromosomal arms. As previously spotted, in WT plants the telomere proximal regions frequently interacted with each other through long-range interactions ^64–68^. We further observed that ITR-1R and ITR-4L do not particularly associate with each other or with telomeres (Figure S14A). In *2h1* nuclei, with the exception of the regions adjacent to the Nucleolar Organizer Regions (NORs) of chromosome 2 and 4 (SubNOR2 and SubNOR4) that displayed atypical patterns (detailed in Figure S14), interaction frequencies between all sub-telomeric regions were increased (Figure 5D). Furthermore, ITR-1R and 4L also showed increased ITR-ITR and ITR-telomere interaction frequency (Figure 5D, S13C, S14C). Consistent with a reduced number of telomere foci in intact *2h1* nuclei (Figure 4), this observation supports an organizational model in which telomeres tend to coalesce more frequently in the absence of H1.

Last, we examined the topology of ITR loci. In WT plants, both formed large structures resembling Topologically Associating Domains (TADs), which are themselves immersed within highly self-interacting pericentromeric regions (S13A). Interestingly, in *2h1* nuclei, intra-ITR interactions were strongly enhanced (*i.e*., TAD re-enforcement) while the surrounding pericentromeric environments expectedly showed an opposite trend linked to heterochromatin relaxation (Figure 5E, S13C). This observation was supported by comparing distal-to-local ratios (DLR) of interaction frequency that showed clear local drops at each ITR in *2h1* nuclei, hence an increased tendency for interacting only with itself usually interpreted as increased domain compaction (Figure S15). ^72^. Altogether, these observations show that, in contrast to its general role in heterochromatin packing, H1 dampens the local insulation of ITRs from their neighboring environment. Remarkably, the boundaries of these compaction defects in *2h1* nuclei sharply correspond with H3K27me3-enrichment (Figure 5E, S15).

### H1 antagonizes TRB-mediated PRC2 activity at ITRs

With the aim to determine the molecular mechanisms by which H1 selectively represses PRC2 activity at ITRs, we envisioned that Telomere Repeat Binding (TRB) proteins might have a prominent role (Figure 6A). The TRB1, TRB2 and TRB3 founding members of this plant-specific Single-Myb-histone proteins constitute part of the telomere nucleoprotein structure required to maintain telomere length ^73^ (Figure 6B). Their Myb domain has strong affinity to the G-rich strand of *telobox* DNA motifs ^73–75^ and, combined with a coiled-coil domain that associates with the CURLY-LEAF (CLF) and SWINGER (SWN) catalytic subunits of PRC2 ^36,37^, TRBs act as transcriptional regulators of proteincoding genes bearing a *telobox* motif ^76,77^. Interestingly, despite their low protein sequence similarity to H1 histones (14±2%; Figure S16), TRBs display a typical GH1 domain ^19,78^. Hence, we hypothesized that antagonistic chromatin incorporation of the GH1 domains of TRB and H1 proteins might modulate PRC2 recruitment at ITRs.

**Figure 6.**
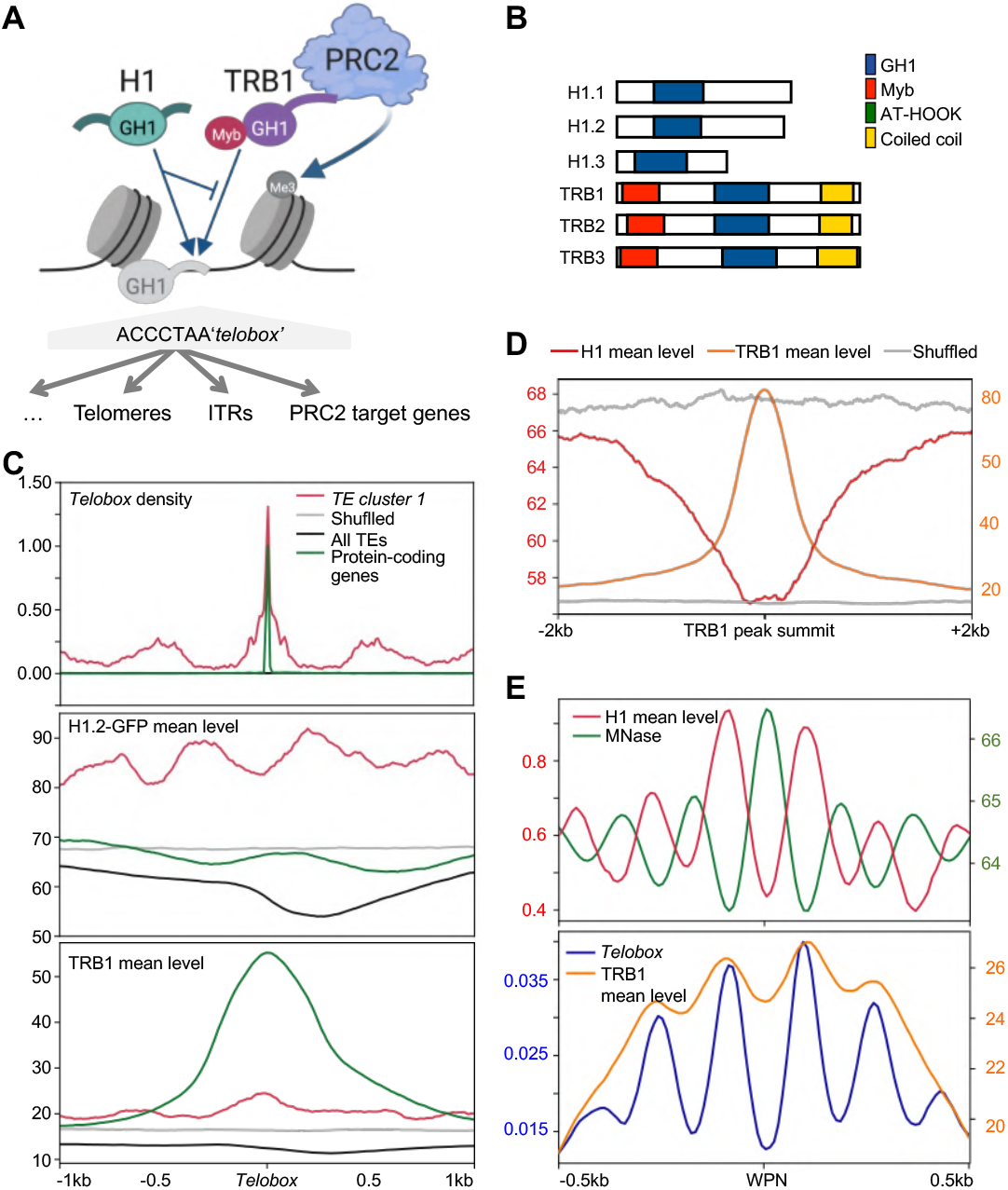
Antagonistic chromatin association of H1 and TRB1 over the genome. **A.** Working model of H1 / TRB1 antagonistic chromatin association at linker DNA-localized *telobox* motifs and its sequence-specific influence on PRC2 recruitment at distinct chromatin regions displaying telomeric repeats. **B.** TRB family members possess an amino-terminal single-Myb domain with sequence specificity for *telobox* motifs, a coiled coil domain enabling their association with PRC2 subunits ^37^, and a central GH1 domain that may trigger competitive binding with H1. **C.** H1 and TRB1 patterns are both influenced by *telobox* positioning and they display an opposite trend at *TE Cluster 1 telobox* motifs. The plots display TRB1 and H1 mean read coverage at all TAIR10 genome *telobox* motifs. **D.** H1 occupancy is reduced at genome loci corresponding to TRB1 peak summits. **E.** H1, TRB1 and *telobox* motifs all tend to associate with DNA linker regions. Genome-wide profiles of H1, TRB1, and *telobox* sequence motifs were plotted over the coordinates of all *Arabidopsis* well-positioned nucleosomes defined by Lyons & Zilberman (2017) ^28^. In (C-D) shuffled controls were prod uced with random permutations of genomic position of the regions of interest.

TRB1 genomic distribution. Analysis of available TRB1 ChIP-seq data ^76^ showed that TRB1 peak summits expectedly correlates with the position of *telobox* motifs located in protein-coding genes. Yet, despite the presence of numerous *telobox* sequences, TRB1 poorly occupies *TE cluster 1* elements (Figure 6C, S17A). Reciprocally, H1 average occupancy is low at TRB1 peaks over the genome (Figure 6D). These observations hint at an antagonistic *cis*enrichment of H1 and TRB1 at chromatin. To better resolve these general patterns and link them to linker DNA positioning, we examined the profiles of H1, TRB1, *telobox* motifs, and nucleosome occupancy around well-positioned nucleosome (WPN) coordinates defined using MNase-seq ^28^. As expected, H1.2-GFP distribution was enriched at DNA linker regions. Surprisingly, this was also the case of *telobox* motif distribution that sharply coincided with regions serving as linker DNA. While TRB1 peaks appeared much broader, their summits are also more pronounced at regions corresponding to linker DNA coordinates (Figure 6E). Hence, if it exists, competitive binding between H1 and TRB proteins likely occurs at linker DNA.

These observations are all compatible with a mechanism by which high H1 occupancy at ITRs prevents TRB1 DNA binding. In consequence, increased access to ITRs in *2h1* mutant plants would facilitate TRB1-mediated PRC2 recruitment. To functionally assess whether this model holds true, we first examined whether GFP-TRB1 accumulates at ITRs in *2h1* plants and then determined the H3K27me3 profile in mutant plants lacking both linker H1 and TRB1, TRB2 and TRB3. To undertake the first experiment, we crossed a *TRB1::GFP-TRB1* line ^76,77^ with *2h1* and revealed GFP-TRB1 genome association by ChIP-seq and ChIP telomere dot-blot. Comparison of GFP-TRB1 chromatin association in WT and *2h1* plants showed a significantly increased association at *TE Cluster 1*, ITR-1R and ITR-4L, and at telomeres (Figure 7, S11B and S18A-B), thereby providing evidence that H1 restricts TRB1 binding to these loci *in vivo*.

**Figure 7.**
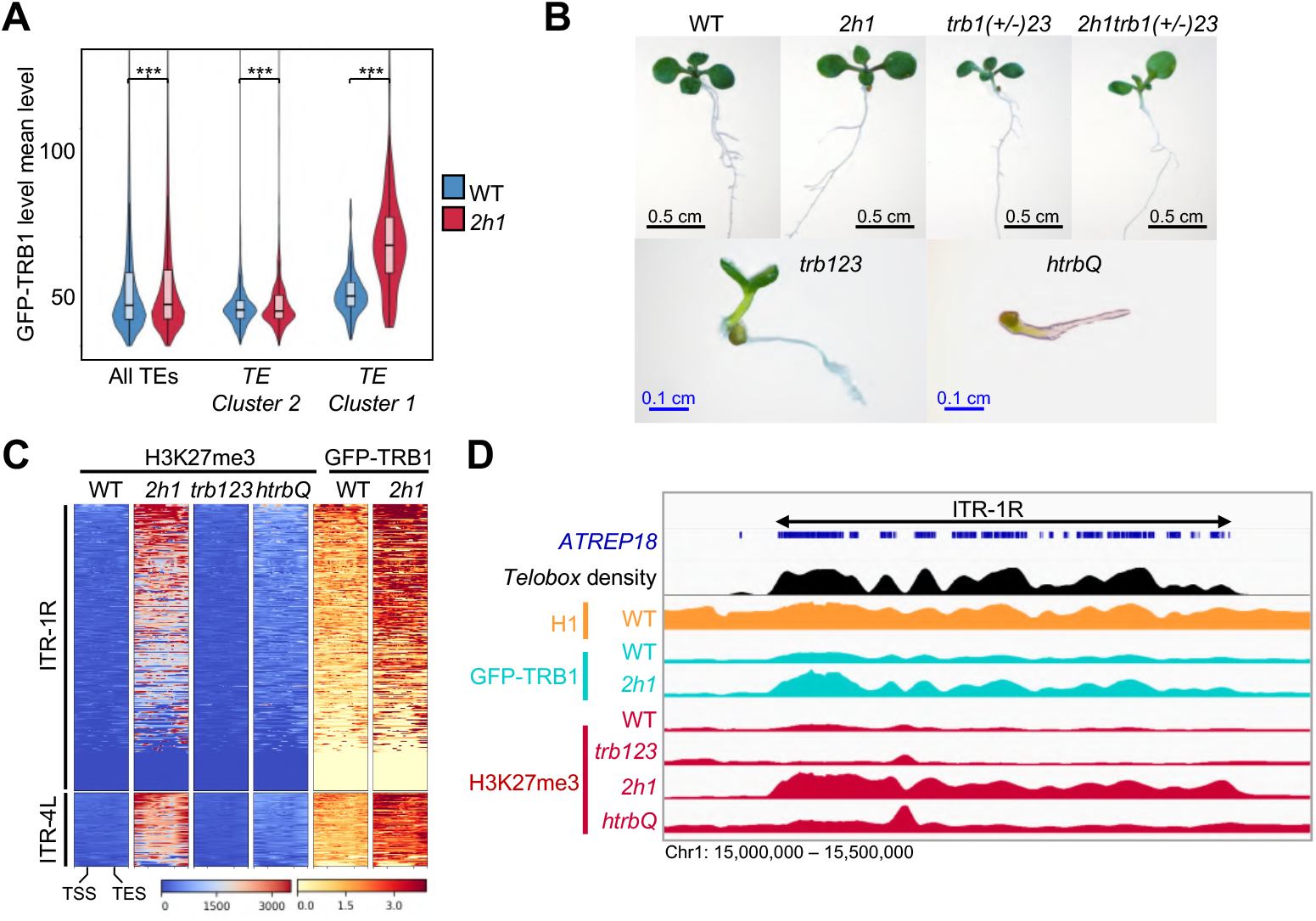
H1 antagonizes TRB1-mediated PRC2 activity at ITRs. **A.** H1 restricts GFP-TRB1 protein association at *TE Cluster 1* elements. The plots show GFP-TRB1 mean normalized coverage in WT and *2h1* seedlings at the indicated repeat categories. *** indicates p-Values <1.94e-05 Wilcoxon signed-rank test. **B.** Homozygous *h1.1h1.2trb1trb2trb3* (*htrbQ*) quintuple mutant seedlings represented 25% of the segregating progeny and displayed strongly altered seedling phenotypes with deficient cotyledon development and slow root growth, indicating that morphogenesis is strongly affected upon combined *H1* and *TRB123* loss-of-function. WT, *2h1* and *trb123* mutant lines have been selected as null F2 segregants from the same cross as the analyzed *htrbQ* plant line. **C.** H1 and TRB proteins are all required for H3K27me3 enrichment at ITR-1R and ITR-4L TEs. TAIR10 annotated repeats located within ITR-1R and ITR-4L coordinates were ranked similarly in all heatmaps. H3K27me3 levels were determined using spike-in normalized ChIP-seq analysis. **D.** Browser view showing that GFP-TRB1 and H3K27me3 enrichment at ITR-1R in *2h1* is lost in *htrbQ* mutant seedlings. Each ChIP series is shown as equally scaled profiles of the indicated genotypes. ChIP-seq and ChIP-Rx data represent the mean of two biological replicates, each.

We then determined whether abolishing simultaneously the expression of linker H1 and TRB1, 2, and 3 proteins impacts PRC2 activity at ITRs. To probe H3K27me3 profiles in *h1.1h1.2trb1trb2trb3* quintuple mutant plants (hereby referred as *htrbQ* for short), we crossed *2h1* double mutant plants to *trb1(+/-)trb2trb3* triple mutant plants propagated as a heterozygous state to accommodate the seedling lethality induced by *TRB123* loss-of-function ^36,37^. Homozygous *htrbQ* mutant seedlings exhibited an aggravated phenotype as compared to the *trb123* triple mutant line (Figure 7B), a synergistic effect presumably reflecting a convergence of *H1* and *TRB123* functions in the regulation of common genes ^79^. Despite the dwarf morphology of the quintuple mutant line, we conducted a ChIP-Rx profiling of H3K27me3 in homozygous WT, *2h1*, *trb123* and *htrbQ* seedlings, all segregating from a single crossed individual. Except at a few loci where H3K27me3 is equally present in WT and *trb123* siblings, in the quintuple mutant seedlings H3K27me3 enrichment was almost completely abolished at ITR-1R and more generally at *TE Cluster 1* elements (Figure 7C-D and S18B-C). Taken together, these analyses demonstrate that H1 occupancy at ITRs antagonizes TRB proteins recruitment, thereby constituting a mechanism preventing invasion of these large chromosome blocks by H3K27me3.

## Discussion

### H1 has a dual impact on H3K27me3 deposition in *Arabidopsis*

We report that *Arabidopsis* H1 is highly enriched at PRC2 target genes where it typically promotes H3K27me3 enrichment and diminishes chromatin accessibility. Contrasting with this general tendency, we also identified an opposite role of H1 in limiting H3K27me3 deposition at interstitial and terminal telomeres as well as at a few genes. This unveiled that, in plants, H1 has a differential effect on H3K27me3 levels over thousands of protein-coding genes on the one hand, and over loci characterized by repeated telomeric motifs on the other hand.

Considering that PRC2 activation is favored at chromatin made of closely neighboring nucleosomes^14^, we postulate that the repertoire of genes losing H3K27me3 upon H1 depletion are those where H1 is required to attain a compaction level enabling efficient PRC2 *cis* activity. Supporting this hypothesis, genes sensitive to H1 for efficient H3K27me3 marking tend to 1) display lower H1 and nucleosome occupancy, 2) be more accessible and 3) be more expressed, than genes unaffected by H1 depletion. In contrast, genes and TEs gaining H3K27me3 upon H1 loss tend to have an elevated nucleosome density and to be weakly accessible while exhibiting sequence signatures potentially triggering different mechanisms of PRC2 regulation such as H1-TRB protein interplays. The large scale on which these antagonistic patterns are observed sheds light on the existence of prominent functional links between H1 and PRC2-based regulation, two main factors in the instruction of DNA accessibility.

### Promoting H3K27me3 enrichment at genes: an evolutionarily conserved function of H1

We identified that H1 has a general role in H3K27me3 deposition at genes, yet, most of the H3K27me3 peaks are still detectable in *2h1* plants. Hence, in agreement with the subtle phenotypes of *h1* mutant plants, H1 is likely not mandatory for the nucleation of PRC2 activity but rather for H3K27me3 maintenance or spreading in *Arabidopsis*. In term of chromatin function, H1 depletion results in a global increase in chromatin accessibility at gene bodies but its impact on expression was apparently more related to variations in H3K27me3 marking. Hence, consistent with the functional categories of the misregulated genes in *2h1* plants, part of the defects in gene expression resulting from H1 depletion might result from indirect consequences on PRC2 activity. The recent findings that depletion of H1 variants in mouse cells triggers widespread H3K27me3 loss and misregulation of PRC2-regulated genes, thereby phenocopying loss of EZH2 ^16,17^, suggest that favoring PRC2 activity is an evolutionarily conserved function of H1.

### H1 hinders PRC2 activity at telomeric repeats by preventing local association of TRB proteins

We provide evidence that H1 antagonizes TRB-mediated PRC2 activity at telomeric repeats. Waiting for an assessment of their relative affinity for *telobox* elements in a chromatin context, H1 / TRB1 proteins antagonistic association along the genome plausibly results from competitive DNA binding of their respective GH1 protein domains. Firstly, chromatin incorporation of H1 and TRB1 is negatively correlated at a genome-wide scale. Secondly, analysis of nucleosome positioning showed that *telobox* motifs are preferentially situated in linker DNA where TRB1 association is also pronounced; so that competition with H1 can occur on linker DNA. Thirdly, profiling of TRB1 chromatin association in *2h1* plants showed that TRB1 ectopically invades ITRs and other *telobox*-rich elements upon H1 loss. These observations reveal that, in wild-type plants, elevated H1 incorporation limits TRB1 enrichment and/or accessibility on these loci despite the presence of repeated *telobox* motifs for which the TRB1 Myb domain has strong affinity ^74,76^.

H3K27me3 profiling in quintuple *2h1trb123* seedlings showed that TRB123 proteins are required for H1-mediated repression of PRC2 activity at telomeric repeats, thereby demonstrating a functional framework in which repression of H3K27me3 deposition at telomeric repeats relies on H1 preventing local association of PRC2-associated TRB proteins. Future studies will determine whether other chromatin modifiers influencing H3K27me3 are implicated. The latter possibility cannot be discarded as, for example, the PRC1 subunit LIKE-HETEROCHROMATIN 1 (LHP1) acting as a chromatin reader of H3K27me3 in *Arabidopsis* ^81^, prevents TRB1 enrichment at PRC2 target genes displaying *telobox* motifs^77^. The outstanding genome-wide pattern of *telobox* positioning in linker DNA also suggests a capacity of this sequence motif to influence chromatin organization, possibly by repelling nucleosomes.

### A new role for H1 on telomeric chromatin structure

Owing to their repetitive nature ^82–87^, the chromatin composition and organization of plant telomeres has long remained enigmatic ^88,89^. ChIP dot-blot analyses indicated a dominance of H3K9me2 over H3K27me3 histone marks ^62,84,90^. Using ChIP dot-blot and *in situ* immunolocalization with telomeric probes we showed that H1 moderates by a 2-to-4 fold the accumulation of H3K27me3 at telomeres. A limitation of our study is that we could not assess the precise distribution of HK27me3 enrichment along each telomere. In agreement with the mosaic chromatin status of telomeres in other organisms^91^, *Arabidopsis* telomeres are thought to be made of segments with distinct nucleosome repeat length (NRL). With average length of 150 bp ^92^, this is much shorter than the 189 bp estimated for H1-rich TEs ^23^. Considering that H1 protects about 20 bp of DNA *in vitro* ^93^, such a small DNA linker size is seemingly incompatible with H1 incorporation into telomere chromatin. For instance, H1 has been proposed to be underrepresented at telomeres in plants ^92,94^ as it is in mammals ^88,95–97^. This could explain the short NRL of *Arabidopsis* and human telomeres ^92,98^ that, long after being suspected ^99^, have recently been re-constructed as a H1-free state columnar organization ^100^. In conclusion, the existence of distinct chromatin states at *Arabidopsis* telomeres needs to be explored in more detail to establish whether the H1-mediated repression of PRC2 activity is a global property of telomeres or rather impacts a few segments.

### H1 has a profound influence on the *Arabidopsis* 3D genome topology

Using Hi-C we identified a reduced frequency of chromatin interactions within and among the pericentromeres in *2h1* nuclei. This is a typical feature of *Arabidopsis* mutants affecting chromocenter formation ^64,65,67^ or when chromocenters get disrupted in response to environmental stress ^68^. These analyses refine the recent observation that chromocenter formation is impaired in *2h1* nuclei ^23,26,35^, a defect that commonly reflects the spatial dispersion of pericentromeres within the nuclear space ^63^. They also shed light on a complex picture in which ITR-1R and 4L embedded within the pericentromeres of chromosomes 1 and 4 escape the surrounding relaxation of heterochromatin induced by H1 depletion and organize themselves as TAD-like structures. In *2h1* nuclei, H3K27me3 invasion at ITRs might underlie the maintenance of compacted and poorly accessible chromatin while neighboring heterochromatic regions tend to become more accessible. It is noteworthy that, in the absence of CTCF and of obvious related 3D structures, *Arabidopsis* is thought to lack proper TADs ^101–103^, hence H1 regulation of ITR insulation represents a new regulatory function of *Arabidopsis* genome topology.

We also report that H1 depletion leads to a reduction in the number of telomeric foci and of their proportion near the nucleolus. This suggested that *2h1* mutants are impaired in telomere spatial individualization, which is indeed supported in our Hi-C analyses by more frequent inter-chromosomal interactions between telomere proximal regions. As the preferential positioning of telomeres around the nucleolus and centromeres near the nuclear periphery is an important organizing principle of *Arabidopsis* chromosome sub-nuclear positioning^63^ and topology^69^, H1 therefore appears to be a crucial determinant of *Arabidopsis* nuclear organization.

Both PRC1 and PRC2 participate in defining *Arabidopsis* genome topology ^64,104^, and H3K27me3 is favored among long-distance interacting gene promoters ^66^. This led to the proposal that, as in animals, this mark could contribute to shape chromosomal organization in *Arabidopsis*, possibly through the formation of *Polycomb* subnuclear bodies ^66,105^. Here we mostly focused on large structural components of the genome, such as telomeres, pericentromeres and ITR regions. In mammals, H1 depletion not only triggers higher-order changes in chromatin compartmentation ^16,17^, but also extensive topological changes of gene-rich and transcribed regions ^18^. Future studies will determine to which extent the impact of H1 on the H3K27me3 landscape contributes to define *Arabidopsis* genome topology.

### H1 as a modulator of H3K27me3 epigenome homeostasis

In *Neurospora crassa*, artificial introduction of an (TTAGGG)_17_ telomere repeats array at interstitial sites was shown to trigger the formation of a large H3K27me2/3-rich chromosome domain ^106^. Followed by our study, this illustrates the intrinsic attractiveness of telomeric motifs for H3K27me3 deposition in multiple organisms. With several thousands of telomeric motifs altogether covering ^~^430 kb, ITRs represent at least twice the cumulated length of all telomeres in *Arabidopsis* diploid nuclei, thereby forming immense reservoirs of PRC2 targets. Our findings led us to hypothesize that H1-mediated repression of PRC2 activity at these scaffolding domains serves as a safeguard to avoid the formation of gigantic H3K27me3-rich blocks in both pericentromeric and telomeric regions, which can be detrimental not only for chromosome folding but could also be on a scale tethering PRC2 complexes away from protein-coding genes. In other terms, balancing PRC2 activity between protein-coding genes and telomeric repeats, H1 protein regulation may represent an important modulator of epigenome homeostasis during development.

## Methods

### Plant lines and growth conditions

The *h1.1 h1.2* (*2h1*) *Arabidopsis* mutant line and the transgenic *pH1.2::H1.2-GFP* line have already been described ^22^ and were kindly provided by Dr. Kinga Rutowicz (University of Zurich, Switzerland). The *2h1/TRB1::GFP-TRB1* transgenic line was obtained upon manual crossing of the *2h1* and *TRB1::GFP-TRB1* line described previously in^76,77^. The *trb123* triple mutant line was produced by crossing a *trb1trb2* double homozygous line derived from a cross between *trb1* (Salk_001540) and *trb2* (Flag_242F11) mutant alleles with the double homozygous *trb2trb3* mutant derived from a cross between *trb2* (Flag_242F11) with *trb3* (Salk_134641). Seeds were surface-sterilized, plated on half strength Murashige and Skoog (MS) medium with 0.9% agar and 0.5% sugar, and cultivated under long-day (16h/8h) at 23/19°C light/dark photoperiod (100 μmol.m^-2^.s^-1^) for 5 days unless otherwise stated. Cotyledons, when used, were manually dissected under a stereomicroscope.

### Immuno-FISH

After fixation in 4% paraformaldehyde in 1X PME, cotyledons of 7-day-old seedlings were chopped directly in 1% cellulase, 1% pectolyase, and 0.5% cytohelicase in 1X PME, and incubated 15 min. Nucleus suspensions were transferred to poly-Lysine-coated slides. One volume of 1% lipsol in 1X PME was added to the mixture and spread on the slide. Then, 1 volume of 4% PFA in 1X PME was added and slides were dried. Immunodetection and FISH were conducted as described previously ^78^ using the following antibodies: rabbit H3K27me3 (#07-449 - Merck) diluted 1:200, Goat biotin anti Rabbit IgG (#65-6140 - ThermoFisher) 1:500, mouse anti-digoxigenin (#11333062910 -ROCHE) 1:125, rat anti-mouse FITC (#rmg101 - Invitrogen) at 1:500 at 1:100, mouse Cy3 anti-biotin antibody (#C5585 - Sigma) at 1:1000. Acquisitions were performed on a structured illumination (pseudo-confocal) imaging system (ApoTome AxioImager M2; Zeiss) and processed using a deconvolution module (regularized inverse filter algorithm). The colocalization was analyzed via the colocalization module of the ZEN software using the uncollapsed Z-stack files. To test for signal colocalization, the range of Pearson correlation coefficient of H3K27m3 vs telomeric FISH signals were calculated with the colocalization module of the ZEN software using Z-stack files. Foci with coefficients superior to 0.5 were considered as being colocalized.

### ATAC-seq

Nuclei were isolated from 200 cotyledons of 5-day-old seedlings and purified using a two-layer Percoll gradient at 3000 g before staining with 0.5 μM DAPI and sorting by FACS according to their ploidy levels using a MoFlo Astrios EQ Cell Sorter (Beckman Culture) in PuraFlow sheath fluid (Beckman Coulter) at 25 psi (pounds per square inch), with a 100-micron nozzle. We performed sorting with ^~^43 kHz drop drive frequency, plates voltage of 4000-4500 V and an amplitude of 30-50 V. Sorting was performed in purity mode. For each sample, 20000 sorted 4C nuclei were collected separately in PBS buffer and centrifuged at 3,000 g at 4 °C for 5 min. The nuclei were re-suspended in 20 μl of Tn5 transposase reaction buffer (Illumina). After tagmentation, DNA was purified using the MinElute PCR Purification Kit (Qiagen) and amplified with Nextera DNA Library Prep index oligonucleotides (Illumina). A size selection was performed with AMPure^®^ XP beads (Beckman Coulter) to collect library molecules longer than 150 bp. DNA libraries were sequenced by Beijing Genomics Institute (BGI Group, Hong-Kong) using the DNA Nanoballs (DNB^™^) DNBseq in a 65 bp paired-end mode.

### *In situ* Hi-C

Hi-C was performed as in Grob *et al* (2014) ^65^ with downscaling using seedlings crosslinked in 10 mM potassium phosphate pH 7.0, 50 mM NaCl, 0.1 M sucrose with 4 % (v/v) formaldehyde. Crosslinking was stopped by transferring seedlings to 30ml of 0.15 M glycine. After rinsing and dissection, 1000 cotyledons were flash-frozen in liquid nitrogen and ground using a Tissue Lyser (Qiagen). All sample were adjusted to 4 ml using NIB buffer (20 mM Hepes pH7.8, 0.25 M sucrose, 1 mM MgCl2, 0.5 mM KCl, 40 % v/v glycerol, 1 % Triton X-100) and homogenized on ice using a Dounce homogenizer. Nuclei were pelleted by centrifugation and resuspended in the DpnII digestion buffer (10 mM MgCl2, 1 mM DTT, 100 mM NaCl, 50 mM Bis-Tris-HCl, pH 6.0) before adding SDS to a final concentration of 0.5 % (v/v). SDS was quenched by adding 2% Triton X-100. DpnII (200 u) was added to each sample for over-night digestion at 37 °C. dATP, dTTP, dGTP, biotinylated dCTP and 12 μl DNA Polymerase I (Large Klenow fragment) were added before incubation for 45 min at 37 °C. A total of 50 unit of T4 DNA ligase along with 7 μl of 20 ng/μl of BSA (Biolabs) and 7 μl of 100 mM ATP were added to reach a final volume of 700μl. Samples were incubated for 4h at 16°C with constant shaking at 300rpm. After over-night reverse crosslinking at 65°C and protein digestion with 5 μl of 10 mg/μl proteinase K, DNA was extracted by phenol/chloroform purification and ethanol precipitation before resuspension in 100μL of 0.1X TE buffer. Biotin was removed from the unligated fragment using T4 DNA polymerase exonuclease activity. After biotin removal, the samples were purified using AMPure beads with a 1.6X ratio. DNA was fragmented using a Covaris M220 sonicator (peak power 75W, duty factor 20, cycles per burst 200, duration 150 s). Hi-C libraries were prepared using KAPA LTP Library Preparation Kit (Roche) ^65^ with 12 amplification cycles. PCR products were purified using AMPure beads (ratio 1.85X). Libraries were analyzed using a Qubit fluorometer (ThermoFisher) and a TAPE Station (Agilent) before sequencing in a 75 bp PE mode using a DNB-seq platform at the Beijing Genomics Institute (BGI Group; Honk Kong).

### RNA-seq

Seedlings grown in long days were fixed in 100% cold acetone under vacuum for 10 min. Cotyledons from 100 plants were dissected and ground in 2 ml tubes using a Tissue Lyser (Qiagen) for 1 min 30 sec at 30 Hz before RNA extraction using the RNeasy micro kit (Qiagen). RNA was sequenced using the DNBseq platform at the Beijing Genomics Institute (BGI Group) in a 100 bp paired-end mode. For raw data processing, sequencing adaptors were removed from raw reads with trim_galore! v2.10 (https://github.com/FelixKrueger/TrimGalore). Reads were mapped onto combined TAIR10 genome using STAR version 2.7.3a ^107^ with the following parameters “--alignIntronMin 20 --alignIntronMax 100000 --outFilterMultimapNmax 20 --outMultimapperOrder Random --outFilterMismatchNmax 8 --outSAMtype BAM SortedByCoordinate --outSAMmultNmax 1 --alignMatesGapMax 100000”. Gene raw counts were scored using the htseq-count tool from the HTSeq suite version 0.11.3 ^108^ and analyzed with the DESeq2 package ^80^ to calculate Log_2_-fold change and to identify differentially expressed genes (p-value < 0.01). TPM (Transcripts per Million) were retrieved by dividing the counts over each gene by its length in kb and the resulting RPK was divided by the total read counts in the sample (in millions). Mean TPM values between two biological replicates were used for subsequent analyses. To draw metagene plots, genes were grouped into expressed or not and expressed genes split into four quantiles of expression with the function ntile() of the R package dplyr (https://CRAN.R-project.org/package=dplyr).

### H1.2-GFP, GFP-TRB1 and H3 ChIP-seq experiments

H1.2-GFP and parallel H3 profiling were conducted as in Fiorucci *et al* (2019) ^109^ sonicating chromatin to reach mono/di-nucleosome fragment sizes. WT Col-0 or *pH1.2::H1.2-GFP* seedlings were crosslinked for 15 min using 1 % formaldehyde. After dissection, 400 cotyledons were ground in 2 ml tubes using a Tissue Lyser (Qiagen) for 2 x 1 min at 30 Hz. After resuspension in 100 μl Nuclei Lysis Buffer 0.1 %SDS, the samples were flash frozen in liquid nitrogen and chromatin was sheared using a S220 Focused-ultrasonicator (Covaris) for 17 min at peak power 105 W, duty factor 5%, 200 cycles per burst, to get fragment sizes between 75 and 300 bp. Immunoprecipitation was performed on 150 μg of chromatin quantified using the Pierce^™^ BCA Protein Assay Kit (Thermo Fisher Scientific) with 60 μl of Protein-A/G Dynabeads and 3.5 μl of anti-GFP (Thermo Fisher #A11122) for H1.2-GFP and mock (WT) sample or anti-H3 (Abcam #Ab1791) for H3 IPs. Immunoprecipitated DNA was subjected to library preparation using the TruSeq^®^ ChIP Sample Preparation Kit (Illumina) and sequenced using a NextSeq 500 system or DNBSEQ-G400 in a single-end 50 bp mode (Genewiz, USA; Fasteris, Switzerland and DNBseq BGI, Hong-Kong).

### H3K27me3 ChIP-Rx

ChIP-Rx of WT and *2h1* plants corresponding to Figures 1–6 and of WT, *2h1*, *trb123* and *htrbQ* plants corresponding to Figure 7 were performed using anti-H3K27me3 #07-449 (Millipore) and #C15410069 (Diagenode), respectively. Both ChIP-Rx series were conducted as in Nassrallah *et al* (2018) ^45^ using two biological replicates of 8-day-old WT and *2h1* seedlings. For each biological replicate, two independent IPs were carried out using 120 μg of *Arabidopsis* chromatin mixed with 3 % of Drosophila chromatin quantified using the Pierce^™^ BCA Protein Assay Kit (Thermo Fisher Scientific). DNA samples eluted and purified from the two technical replicates were pooled before library preparation (Illumina TruSeq ChIP) and sequencing (Illumina NextSeq 500, 1×50bp or DNBSEQ-G400, 1×50bp) of all input and IP samples by Fasteris (Geneva, Switzerland) and BGI (Hong-Kong), respectively.

### H3K27me3 and H3 ChIP-blot analyses

Anti-H3K27me3 (Millipore, #07-449 antibody) and anti-H3 (Abcam #Ab1791 antibody) ChIPs were conducted using 2 g of tissue. Pellets of both inputs (20%) and immunoprecipitated DNA were resuspended in 40 μl of TE, pH 8.0 and analyzed through dot-blot hybridization using a radioactively labeled telomeric probe synthesized by non-template PCR ^62,110^. ITRs contribution to the hybridization signal was minimized using high stringency hybridization as detailed in ^62^.

### Hi-C bioinformatics

Mapping of Hi-C reads was performed using the Hi-C Pro pipeline ^111^ with default pipeline parameters and merging data from three biological replicates at the end of the pipeline. Data were in visualized using the Juicebox toolsuite ^112^ and represented in Log_10_ scale after SCN normalization ^113^ with Boost-HiC ^114^ setting alpha parameter to 0.2. In Figure S17, we normalized the sequencing depth in each sample and scored the number of reads in each combination of genomic regions using HOMER ^72^. Read counts were further normalized for the bin size and the median value between the three biological replicates was reported. Distal-to-Local [log2] Ratios (DLR) where implemented as described in HOMER ^72^ and adapted to define local interactions between a defined size window (k) and the two surrounding windows as distal regions at 10kb and 100kb for k=2 to k=150 bins and selected for each ITR a windows value corresponding of 3 ITR sizes (1050 kb for ITR-1R and 240 kb for ITR-4L).

### ChIP-seq and ChIP-Rx bioinformatics

For H3K27me3 spike-in normalized ChIP-Rx, raw reads were pre-processed with Trimmomatic v0.36 ^115^ to remove leftover Illumina sequencing adapters. 5’ and 3’ ends with a quality score below 5 (Phred+33) were trimmed and reads shorter than 20 bp after trimming were discarded (trimmomatic-0.36.jar SE -phred33 INPUT.fastq TRIMMED_OUTPUT.fastq ILLUMINACLIP:TruSeq3-SE.fa:2:30:10 LEADING:5 TRAILING:5 MINLEN:20). We aligned the trimmed reads against combined TAIR10 *Arabidopsis thaliana* and *Drosophila melanogaster* (dm6) genomes with Bowtie2v.2.3.2 using the “--very-sensitive” setting. Duplicated reads and reads mapping to regions with aberrant coverage or low sequence complexity defined in ^116^ were discarded with sambamba v0.6.8. ^117^. Peaks of H3K27me3 read density were called using MACS2 ^118^ with the command “macs2 callpeak -f BAM --nomodel -q 0.01 -g 120e6 --bw 300 --verbose 3 --broad”. Only peaks found in both biological replicates and overlapping for at least 10 % were retained for further analyses. Annotation of genes and TEs overlapping with peaks of histone marks H3K27me3, H3K4me3, and H2Bub were identified using bedtools v2.29.2 intersect as for H3K27me3. We scored the number of H3K27me3 reads overlapping with marked genes using bedtools v2.29.2 multicov and analyzed them with the DESeq2 package ^80^ in the R statistical environment v3.6.2 to identify the genes enriched or depleted in H3K27me3 in *2h1* plants (p-value < 0.01). To account for differences in sequencing depth we used the function SizeFactors in DESeq2, applying a scaling factor calculated as in Nassrallah *et al* (2018) ^45^.

For GFP-TRB1, H1.2-GFP and H3 ChIP-seq datasets, raw reads were processed as for H3K27me3. We counted the reads over genes and TEs using bedtools v2.29.2 multicov and converted them in median counts per million, dividing the counts over each gene or TE by its length and by the total counts in the sample and multiplying by 10^6^ to obtain CPMs (Counts per Million reads). Mean read coverage was used in Figure 1A, while the ratio between median value between biological replicates in IP and median value in Input was used for violin-plot analysis of H1.2-GFP in Figure S6B and S17A. To include nucleosomes in close proximity to gene TSS, an upstream region of 250 bp was also considered for the overlap (minimum 150 bp) for H3K27me3, TRB1 and H3K4me3 (GEO datasets given in Additional file 3). H3K27me3 *TE cluster 1* and *TE cluster 2* were identified using Deeptools plotHeatmap using the --kmeans option set at 2. Tracks were visualized using Integrative Genomics Viewer (IGV) version 2.8.0 ^119^. Meta-gene plots and heatmaps were generated from depth-normalized read densities using Deeptools computeMatrix, plotHeatmap, and plotProfile. Violin-plots, histograms and box-plots were drawn using the package ggplot2 v3.2.1 (https://cran.r-project.org/web/packages/ggplot2/) in the R statistical environment. All scripts used will be made publicly available. Shuffled controls were produced with random permutations of genomic position of the regions if interest. The permutations were generated with bedtools v2.29.2 and the command “bedtools shuffle -chromFirst - seed 28776 -chrom”.

### MNase-seq bioinformatics

MNase read density ^28^ was obtained from NCBI GEO under the accession GSE96994. Genomic location of WPNs shared between WT and *2h1* plants were identified as overlapping WPN coordinates between the two genotypes calculated with bedtools v2.29.2 intersect.

### ATAC-seq bioinformatics

Raw ATAC-seq data were treated using the custom-designed ASAP (ATAC-Seq data Analysis Pipeline; https://github.com/akramdi/ASAP) pipeline. Mapping was performed using Bowtie2 v.2.3.2 ^120^ with parameters --very-sensitive -X 2000. Mapped reads with MAPQ<10, duplicate pairs, and reads mapping to the mitochondrial genome as well as regions with aberrant coverage of low sequence complexity defined in ^116^ were filtered out. Concordant read pairs were selected and shifted as previously described by 4 bp ^121^. Peak calling was performed using MACS2 ^118^ using broad mode and the following parameters: --nomodel --shift −50 --extsize 100. Heatmaps and metaplots were produced from depth-normalized read coverage (read per million) using the Deeptools suite ^122^.

### Statistics

Unless stated otherwise, statistical tests were performed with the R package rstatix_0.7.1 (https://CRAN.R-project.org/package=rstatix) using the functions *wilcox_test* and *wilcox_effsize*. All pairwise comparisons between the read coverage in WT and 2h1 over a given set of gene or TEs were tested with Wilcoxon signed rank test for paired samples, using the wilcox_test function with the option “paired = TRUE”. All other comparisons were tested with Wilcoxon rank sum test for independent samples, setting the option “paired = FALSE”.

### DNA sequence analyses

Motifs enriched in gene promoters (−500 bp to +250 bp after the TSS) and in annotated units of *TE cluster 1* elements were identified using MEME version 5.1.1 ^123^. The following options were used for promoters: “-dna -mod anr -revcomp -nmotifs 10 -minw 5 -maxw 9” and for TEs: “-dna -mod anr -nmotifs 10 -minw 5 -maxw 9 -objfun de -neg Araport11_AllTEs.fasta -revcomp -markov_order 0 -maxsites 10000” where Araport11_AllTEs.fasta correspond to the fasta sequence of all TEs annotated in Araport11.

*Telobox* positioning was analyzed using the coordinates described in ^37^ and obtained from https://gbrowse.mpipz.mpg.de/cgi-bin/gbrowse/arabidopsis10_turck_public/?l=telobox;f=save+datafile. *Telobox* repeat numbers were scored over 10-bp non-overlapping bins, smoothed with a 50-bp sliding window and subsequently used to plot *telobox* density.

### Gene ontology analysis

Gene ontology analysis of H3K27me3 differentially marked genes were retrieved using the GO-TermFinder software ^124^ via the Princeton GO-TermFinder interface (http://go.princeton.edu/cgi-bin/GOTermFinder). The REVIGO ^125^ platform was utilized to reduce the number of GO terms and redundant terms were further manually filtered. The Log_10_ p-values of these unique GO terms were then plotted with pheatmap (https://CRAN.R-project.org/package=pheatmap) with no clustering.

### Protein alignment

Protein sequences of H1.1, H1.2, H1.3, TRB1, TRB2 and TRB3 were aligned using T-Coffee (http://tcoffee.crg.cat/apps/tcoffee/do:regular) with default parameters. Pairwise comparison for similarity and identity score were calculated using Ident and Sim tool (https://www.bioinformatics.org/sms2/ident_sim.html).

## Data and materials availability

This study did not generate new unique reagents. All public genomic data used in this study are listed in Additional file 4. Processed ChIP-Rx/ChIP-seq and RNA-seq data are given in the Supplementary Tables 1 and 2, respectively. Raw genome-wide data generated in this study (Hi-C, ATAC-seq, ChIP-RX, ChIP-seq, RNA-seq) are accessible through the GEO Series accession number GSE160414.

## Graphics

were created using Biorender.com

## Author contributions

GT, LW, IB and ClB performed ChIP and ChIP-RX experiments; GT and LW performed ATAC-seq experiments; KA and MF performed telomere dot-blots; IB performed phenotypic analyses; MB contributed to FACS nucleus sorting; SA performed cytological experiments; GT and SG generated the Hi-C datasets. AK and VC developed ATAC-seq bioinformatics tools. LoC, GT, ClB performed RNA-seq, MNase-seq ChIP-seq, and ATAC-seq bioinformatics analyses. LoC, LeC and SG performed Hi-C bioinformatics. GT, ClB, LoC and FB conceived the study. FB, ClB, AC, CeB, ChB, AC, SA, AP, UG, PPS, JF, and SG supervised research. FB wrote the manuscript and all authors edited the manuscript.

## Lead contact

Further information and requests for resources and reagents should be directed to and will be fulfilled by the Lead Contact, Fredy Barneche.

## Acknowledgements

The authors are grateful to Erwann Cailleux (IBENS, Paris, France) and David Latrasse (IPS2, Orsay, France) for technical guidance with ATAC-seq; Nicolas Valentin (I2BC, Gif, France) for assistance with FACS; Magali Charvin (IBENS, Paris, France) for technical assistance with the IBENS plant growth facility, Frédérique Perronet (IBPS, France) for providing Drosophila samples; Kinga Rutowicz (University of Zurich, Switzerland) and Angélique Déléris (IBENS, Paris and I2BC, Gif-Sur-Yvette, France) for sharing unpublished results.

## Supplementary figures

**Figure S1.**
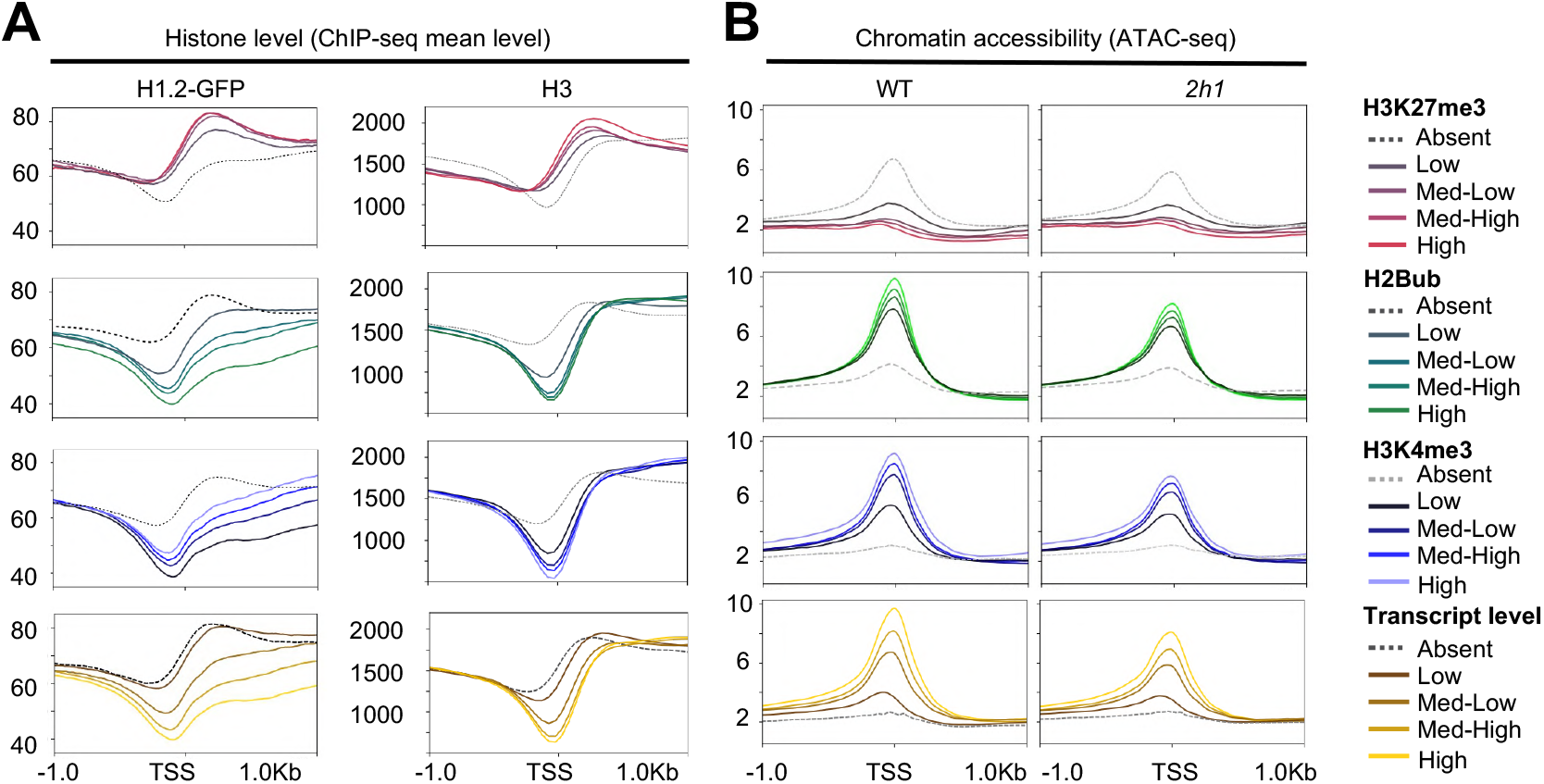
H1.2 distribution and DNA accessibility and their link to gene expression. **A.** H1.2-GFP and H3 levels over the TSS (+/- 1 kp) of genes marked by histone modifications characteristic of PRC2-based repression (H3K27me3; n=7542), transcription initiation (H3K4me3; n=18735), transcription elongation (H2Bub; n=11357) or according to gene expression quartiles (n=27997). Genes with no detectable reads in WT plants were considered as not expressed (n=5894). All ChIPs have been generated in this study except H2Bub and H3K4me3 data given in Additional file 3. **B.** Same analysis as in (A) for ATAC-seq. Mean read coverage is used as a proxy of chromatin accessibility in WT and *2h1* plants.

**Figure S2.**
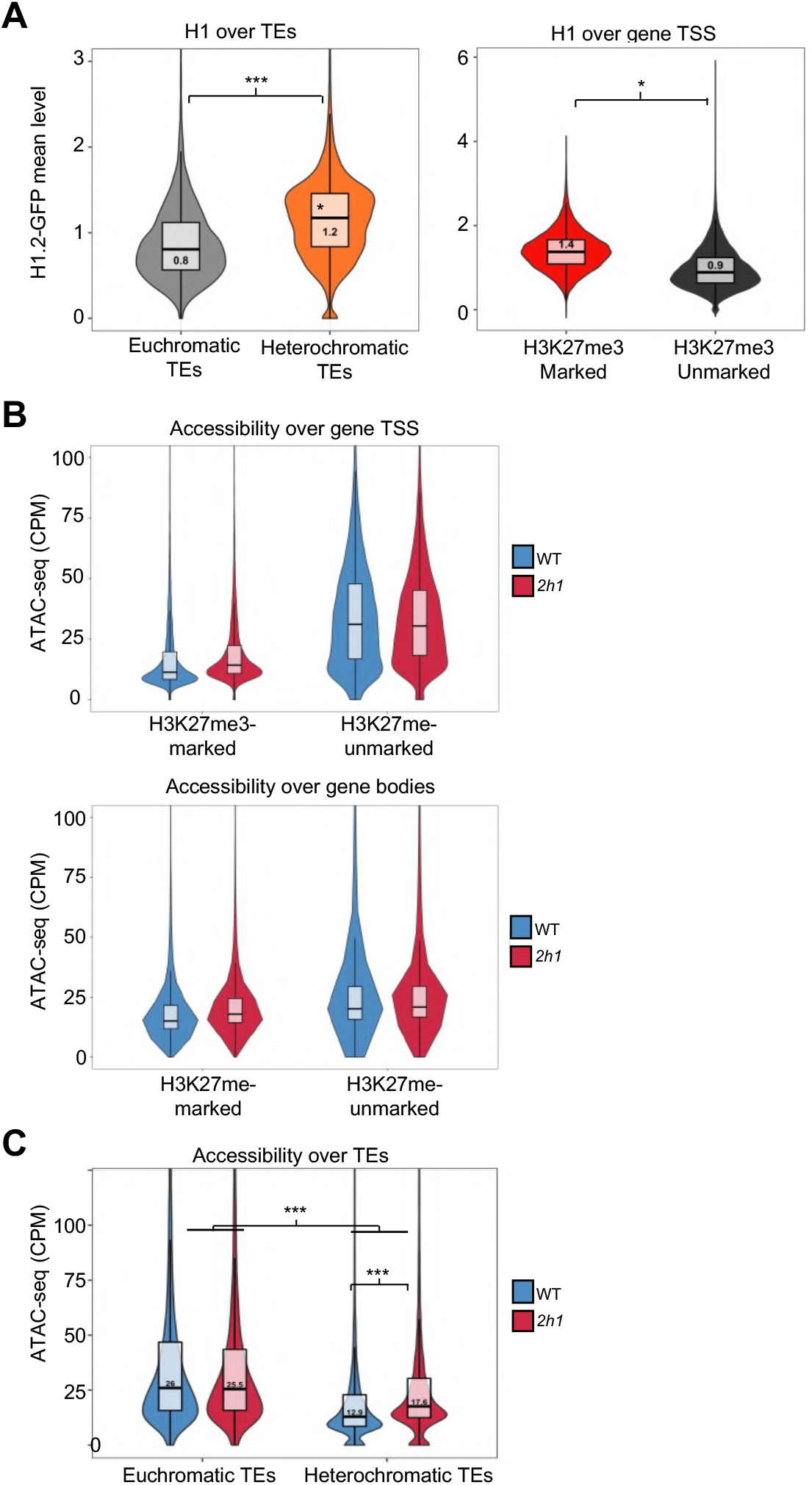
H1.2 distribution and chromatin accessibility properties of TEs and H3K27me3-marked genes. **A.** H1.2-GFP mean level at euchromatic and heterochromatic TEs (mean read coverage). Same analysis than in Figure S1A using TE annotation defined in Bernatavichute et al., (2008). For genes pValue < 2.2e-16; for TEs pValue <1e-308 using a Wilcoxon rank sum test; effect size, moderate. **B.** ATAC-seq read mean density (CPM) over the TSS (+/- 250bp) and full gene body of H3K27me3-marked genes (n=7542) compared to all other annotated protein-coding genes. **C.** Same analysis than (B) for the whole annotations of the corresponding TE sets. Data correspond to the mean of two biological replicates. * and ** indicate a pValue < 10-^122^ and < 10-^308^, respectively, using a Wilcoxon rank test.

**Figure S3.**
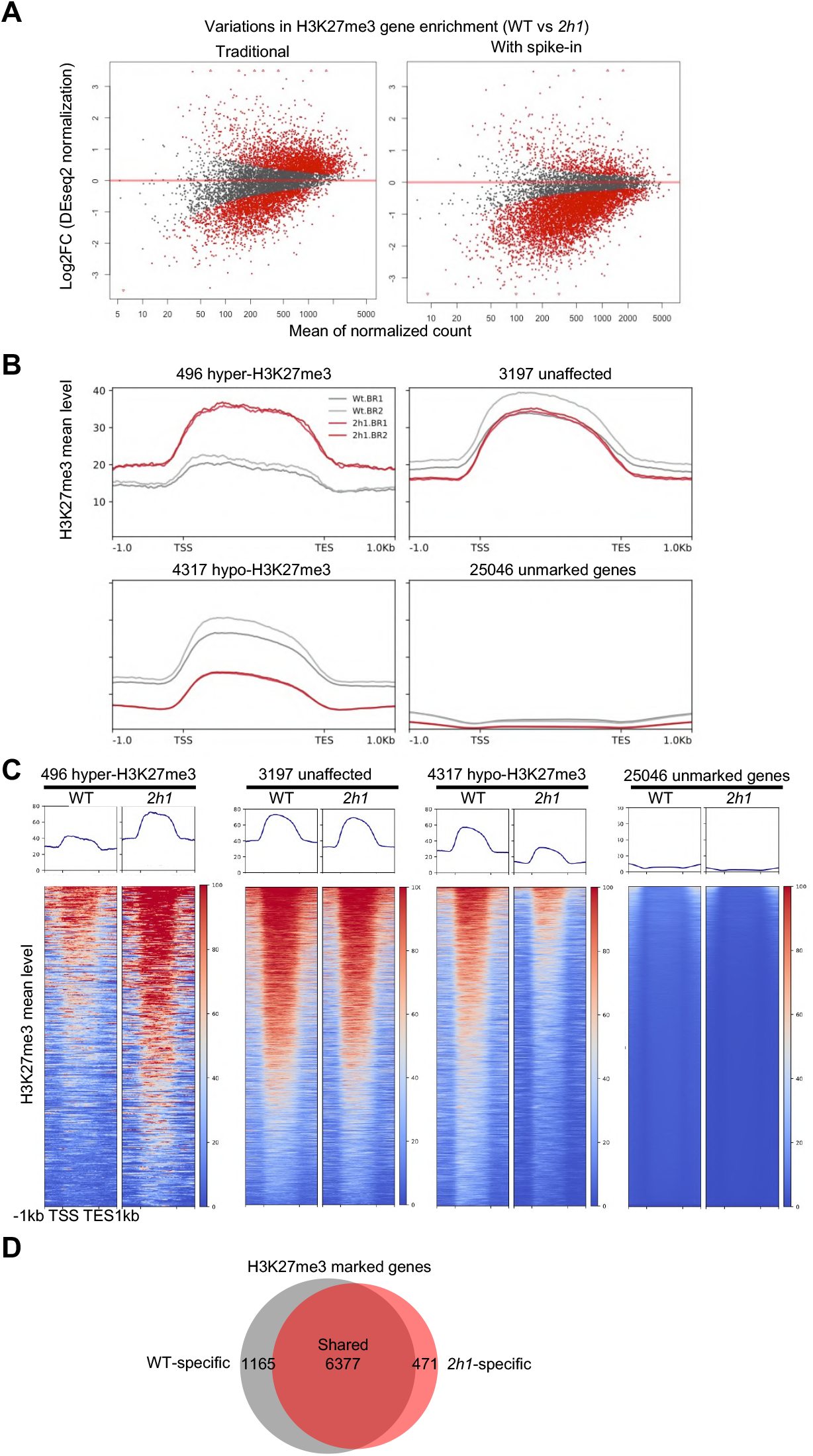
Comparison of the repertoire of genes significantly marked by H3K27me3 in WT and *2h1* plants. **A.** Comparison of the DESeq2 results using either a spike-in normalization factor or DEseq2-based normalization. The plots were drawn using two biological replicates for each sample. **B.** H3K27me3 levels at differentially marked genes (normalized mean read coverage). **C.** Detailed analysis of data presented in Figure 2B. In each cluster genes were ranked according to mean H3K27me3 level. **D.** Number of H3K27me3-marked genes in WT and *2h1* seedlings. Data correspond to the mean of two biological replicates.

**Figure S4.**
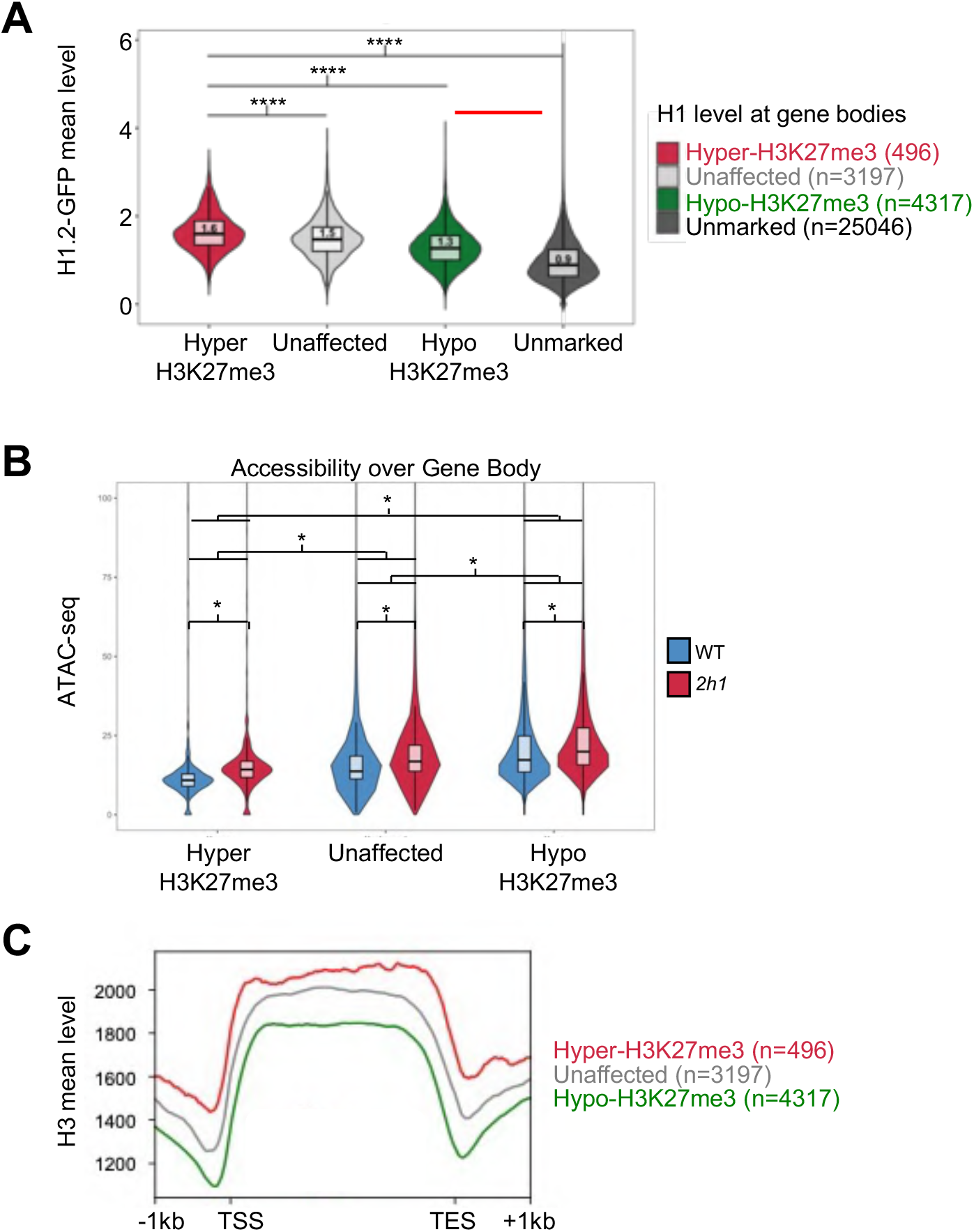
Chromatin properties of H3K27me3 differentially marked genes in *2h1* mutant nuclei. **A.** H1.2-GFP level over the TSS (+/- 250bp) of the gene sets with differential H3K27me3 enrichment in *2h1* plants defined in Figure 2A. **B.** ATAC-seq mean read coverage (CPM) over the whole gene body of the corresponding gene sets. * indicated a pValue < 10-^70^ using a Wilcoxon signed rank test on paired samples. **C.** H3 level over the same gene sets than in (B).

**Figure S5.**
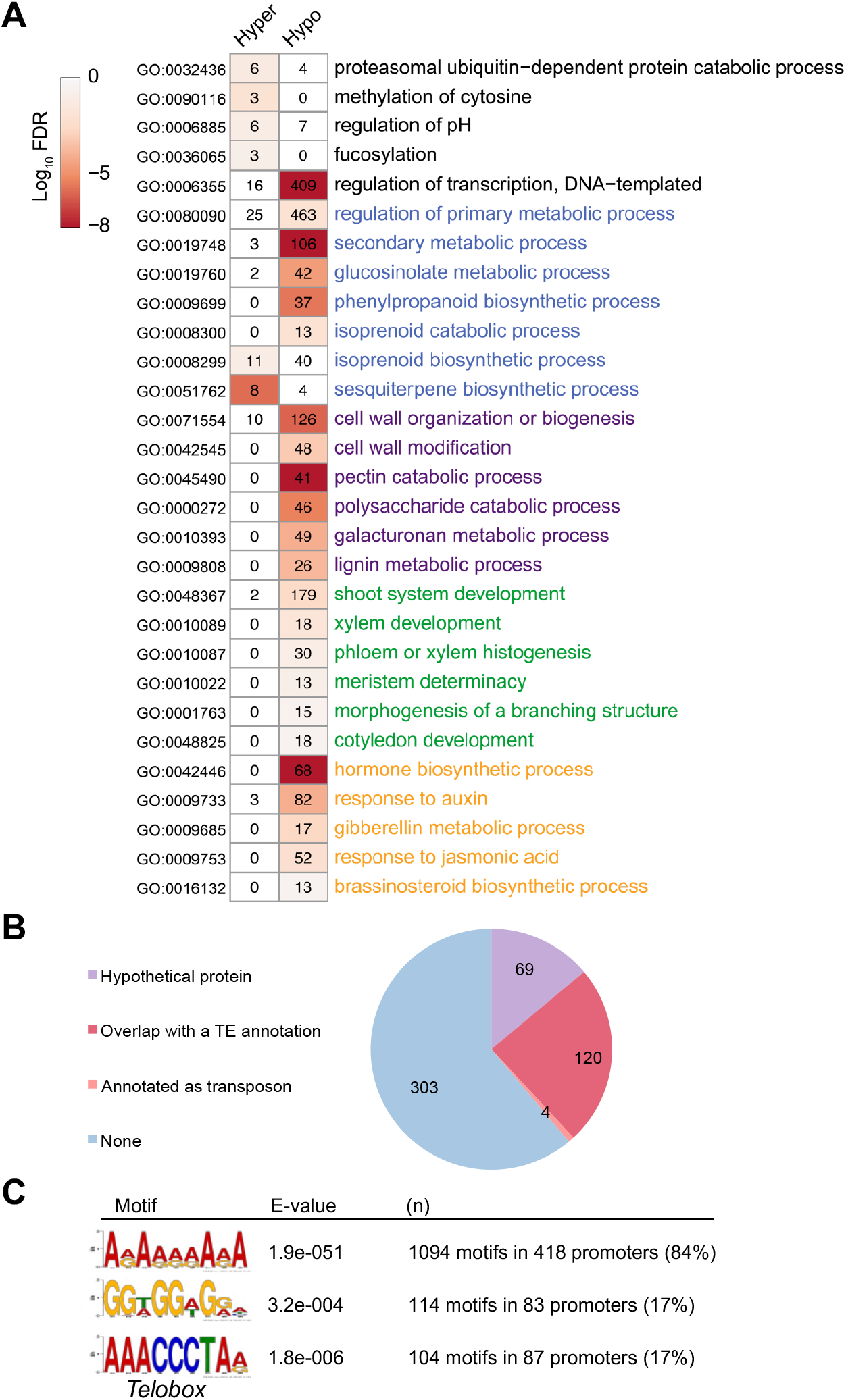
Functional and sequence properties of H3K27me3 differentially marked genes in *2h1* mutant nuclei. **A.** Gene ontology analysis of the genes differentially marked in *2h1* plants. Association to a significantly overrepresented gene function is denoted as a heatmap of false discovery rate (FDR). N=4317 hypo-marked genes; 496 hyper-marked genes. **B.** Number of genes among the 496 hyper-marked genes that either overlap an annotated TE, are annotated as transposons, or are annotated as hypothetical proteins (See Additional file 1 for more details). **C.** Sequence motifs over-represented in the promoters of the 496 H3K27me3 hyper-marked genes in *2h1* plants (Evalue < 1e-220). E-values were calculated against random sequences. The two first motifs could not be matched to any previously known regulatory motif while the 3^rd^ identified motif corresponds to the previously described *telobox* motif (AAACCCTA).

**Figure S6.**
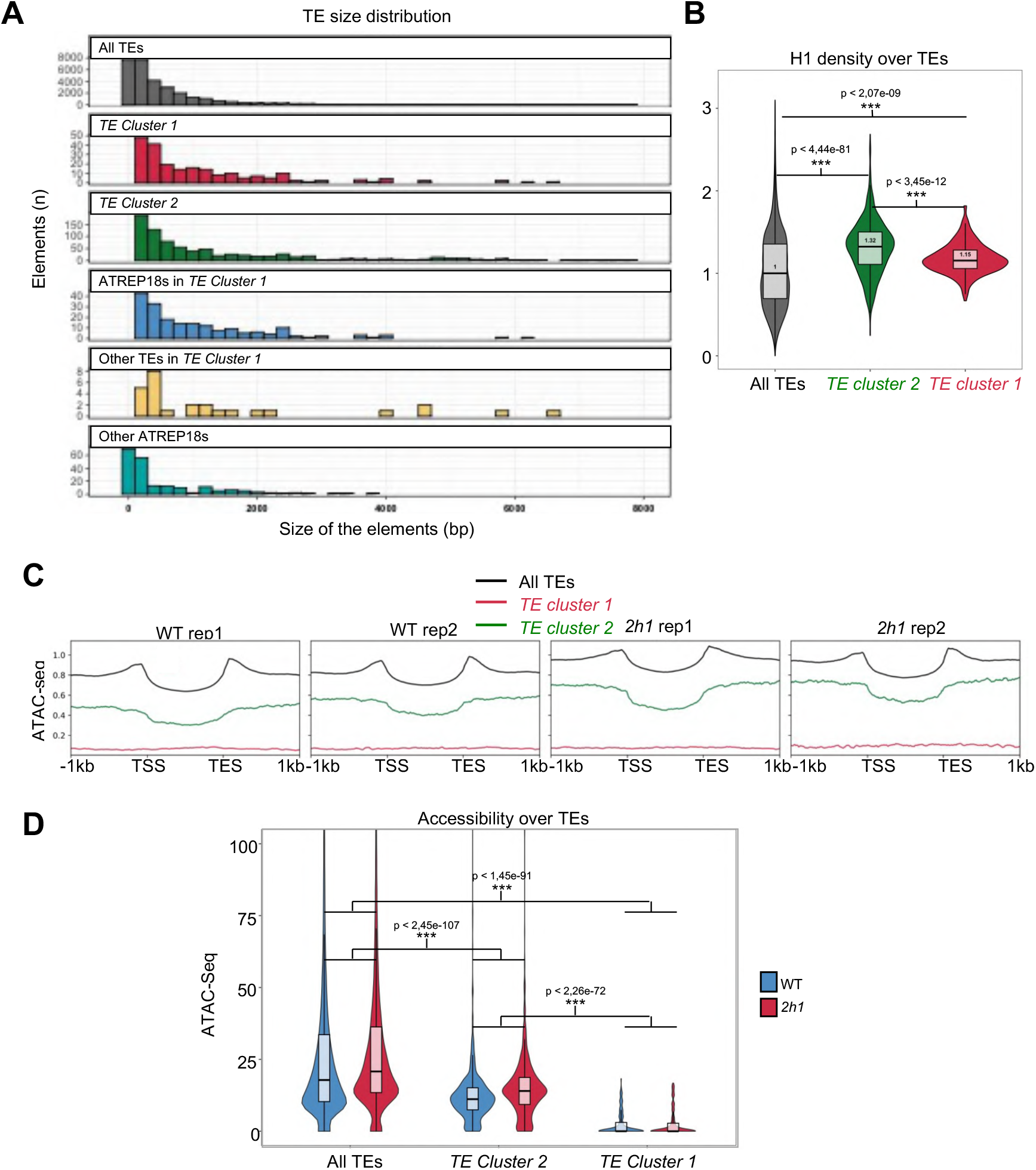
Sequence and chromatin properties of *TE Cluster 1-2* and *ATREP18* elements. **A.** Size distribution of TEs and TE-like repeats belonging to the repertoires defined in Figure 3A. **B.** *TE Cluster 1* is enriched in H1.2-GFP as compared to other TEs (mean read coverage). P-values of differences between the medians assessed using a Wilcoxon rank-sum test test are shown. **C**. Independent biological replicate of ATAC-seq data completing Figure 3D. ATAC-seq mean read coverage of the indicated TE categories in WT and *2h1* nuclei. Profiles obtained for individual ATAC-seq replicates are shown. **D.** ATAC-seq read coverage (CPM) over the whole annotated TE units of corresponding sets. P-values obtained using a Wilcoxon signed rank test on paired samples are given.

**Figure S7.**
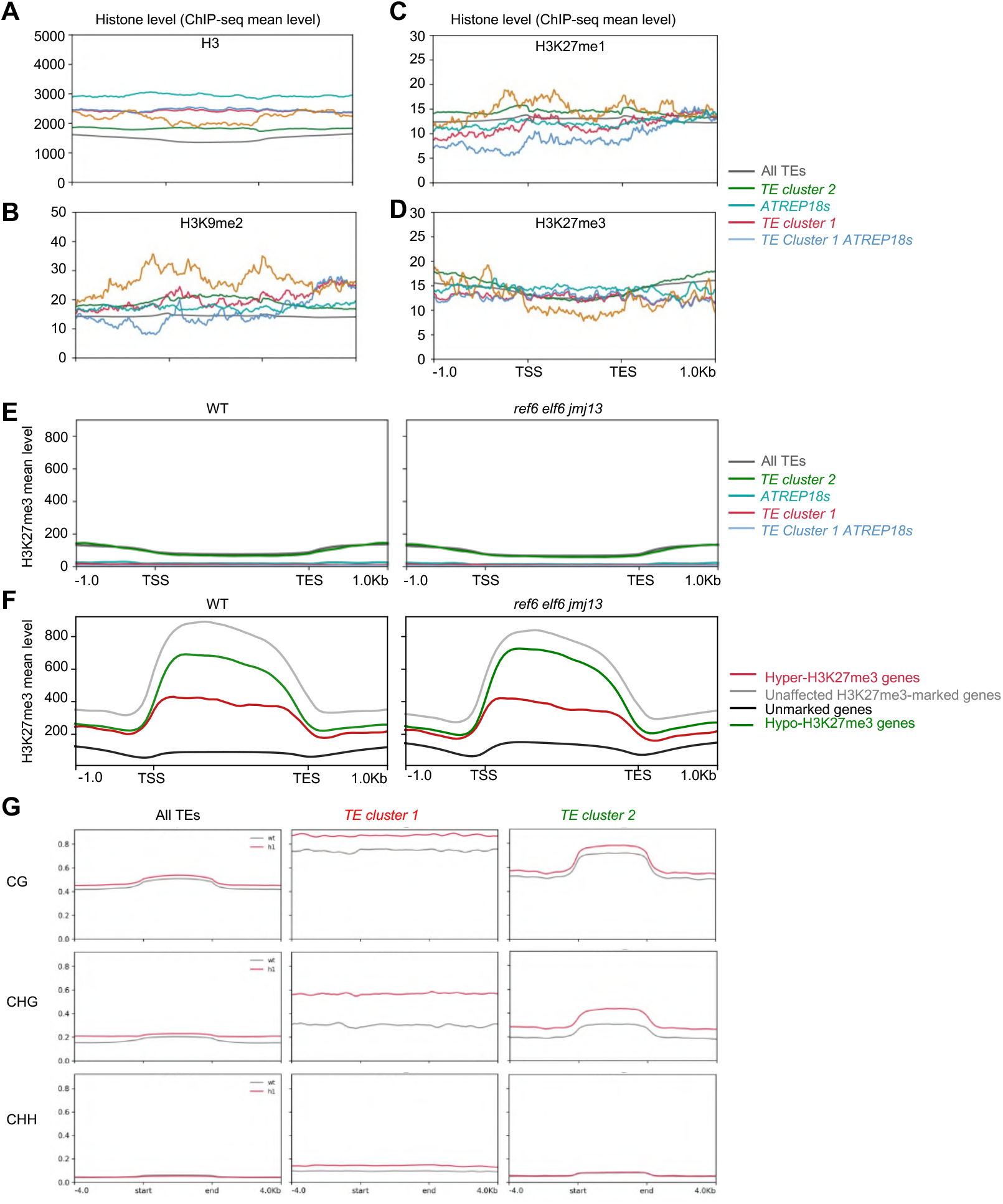
Profile of representative histone marks at *TE clusters 1-2* and *ATREP18* elements. **A-D.** Profiling of H3, H3K9me2, H3K27me1 and H3K27me3 over indicated TE categories. (A) Histone H3 profiling indicates that *ATREP18s* and *TE Cluster1-ATREP18* elements display elevated nucleosome occupancy. This is consistent with the weak accessibility of these elements determined by ATAC-seq analyses. (B) H3K9me2 level is high at *TE Cluster 1* as compared to other TEs. (C) H3K27me1 is not particularly enriched at *TE Cluster 1-2* nor at *ATREP18* elements as compared to other TEs. (D) All TE types investigated in this study, including *ATREP18* elements, display low H3K27me3 levels in WT plants. (A-D) H3K27me3 and H3 but not H3K27me1 and H3K9me2 ChIP-seq have been generated in this study (public data given in Additional file 3). **E.** H3K27me3 profiles of the indicated gene sets in WT and *ref6 elf6 jmj13* triple mutant plants impaired in H3K27me3 demethylation. H3K27me3 data source is given in Additional file 3. **F.** Same analysis than (E) for the indicated gene categories. **G.** CG, CHG and CHH mean methylation at the indicated TE categories in WT and *2h1* mutant seedings. DNA methylation data source is given in Additional file 3.

**Figure S8.**
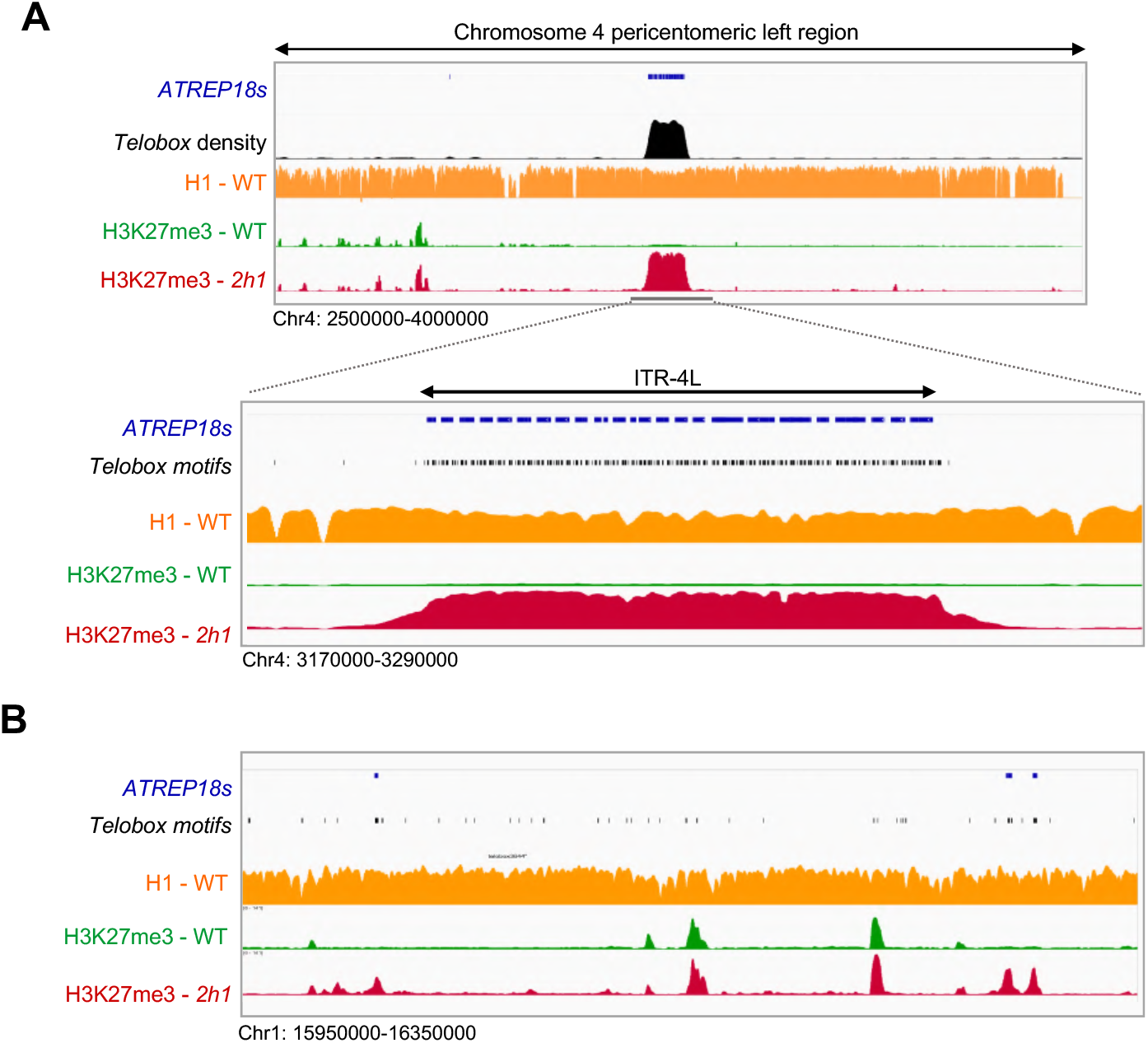
Browser visualization of H3K27me3 profiles at *TE clusters 1-2* and *ATREP18* elements. **A.** Chromosome 4 distribution of *ATREP18* elements, *telobox* motif distribution and H3K27me3 profiles (normalized mean coverage). Top, middle and bottom panels are as in Figure 3G. **B.** Close-up view on interspersed H3K27me3-enriched repeats of *TE cluster 2*, exemplifying a physical correlation between H3K27me3 enrichment and *telobox-* rich domains in *2h1* pericentromeric regions.

**Figure S9.**
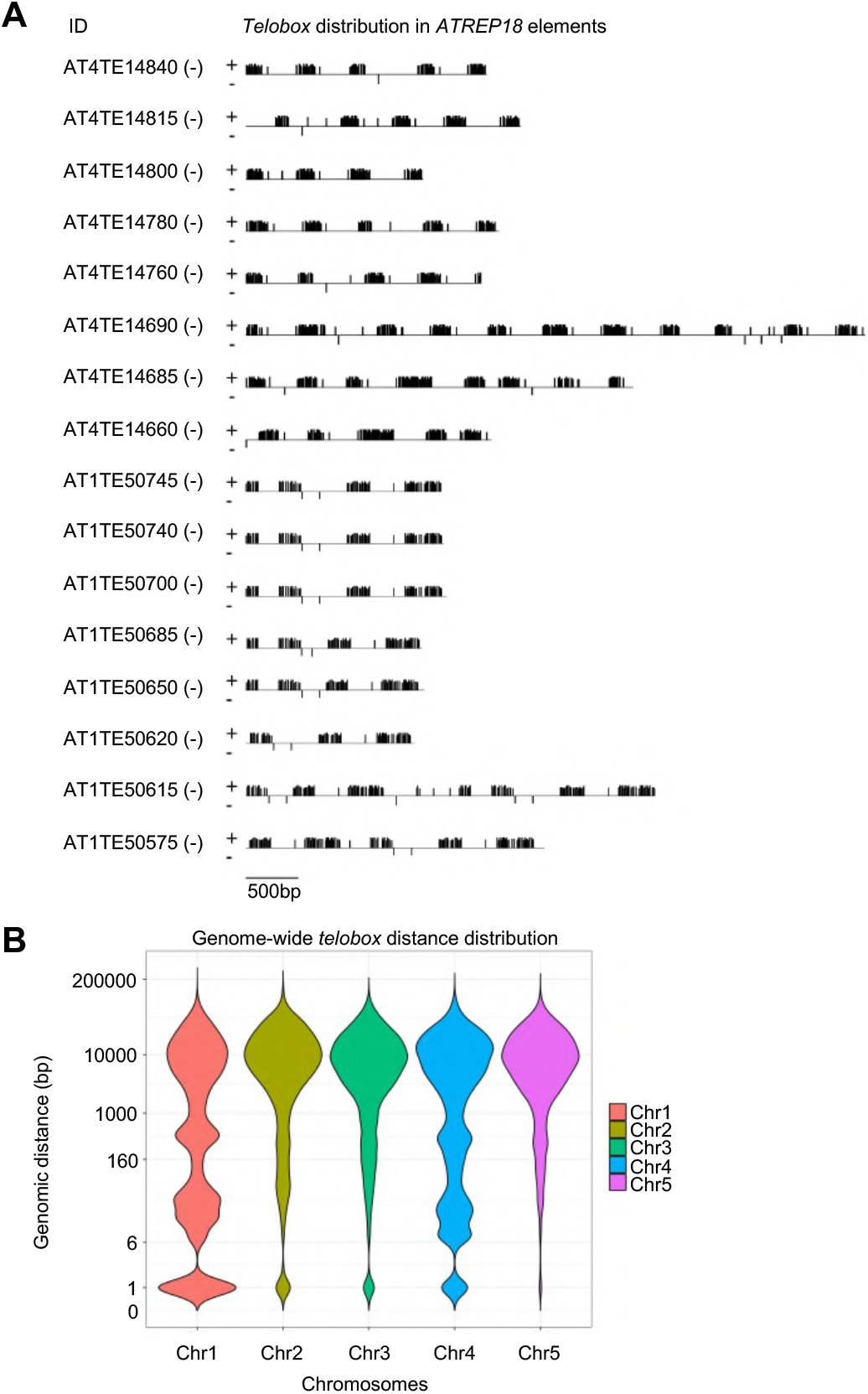
*Telobox* motifs are organized as small clusters in *ATREP18* elements of ITR-1R and ITR-L. **A.** Clustered patterns of *telobox* motif in *ATREP18* elements of *TE Cluster 1*. **B.** Genomic distances between all perfect *telobox* motifs on each chromosome reflecting that ITR organizations in chromosome 1 and 4 constitute important differences as compared to other chromosomes in which interspersed *telobox* motifs are largely prevalent. For example, except at telomeres, chromosome 5 does not display immediately adjacent *telobox* motifs.

**Figure S10.**
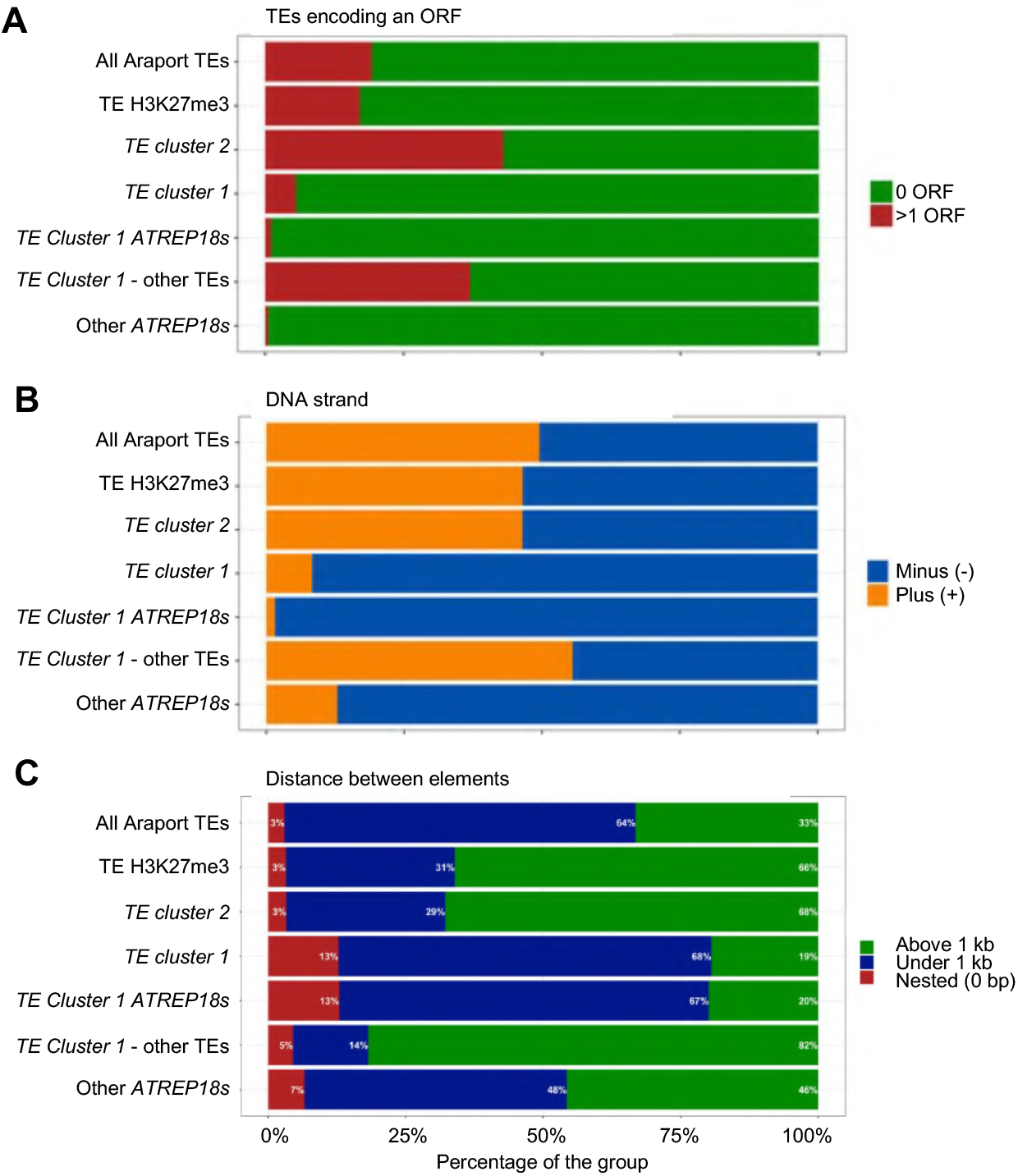
Sequence features of *TE cluster 1-2* elements. **A.** Frequency of open reading regions (ORFs) identified in the indicated TE categories. *TE Cluster 1*, which contains many pericentromeric *ATREP18* elements, rarely encodes ORFs. In contrast, *TE Cluster 2* more frequently encodes ORFs as compared to the ensemble of all TEs. **B.** Strand distribution. *TE Cluster 1* consists mainly of *ATREP18* elements organized in a strand-specific manner. **C.** Distribution of different groups of TEs in three classes of distances: nested (0 base pairs), closely located (under 1 kp) or distantly located (above 1 kb). Compared to the ensemble of all TEs, *TE Cluster 2* elements tend to be dispersed across the genome while, conversely, *TE Cluster 1* elements tend to be located in close proximity. This peculiar distribution is largely due to the overwhelming presence of *ATREP18* elements in *TE Cluster 1* elements in this group (189 out of 216, 87%).

**Figure S11.**
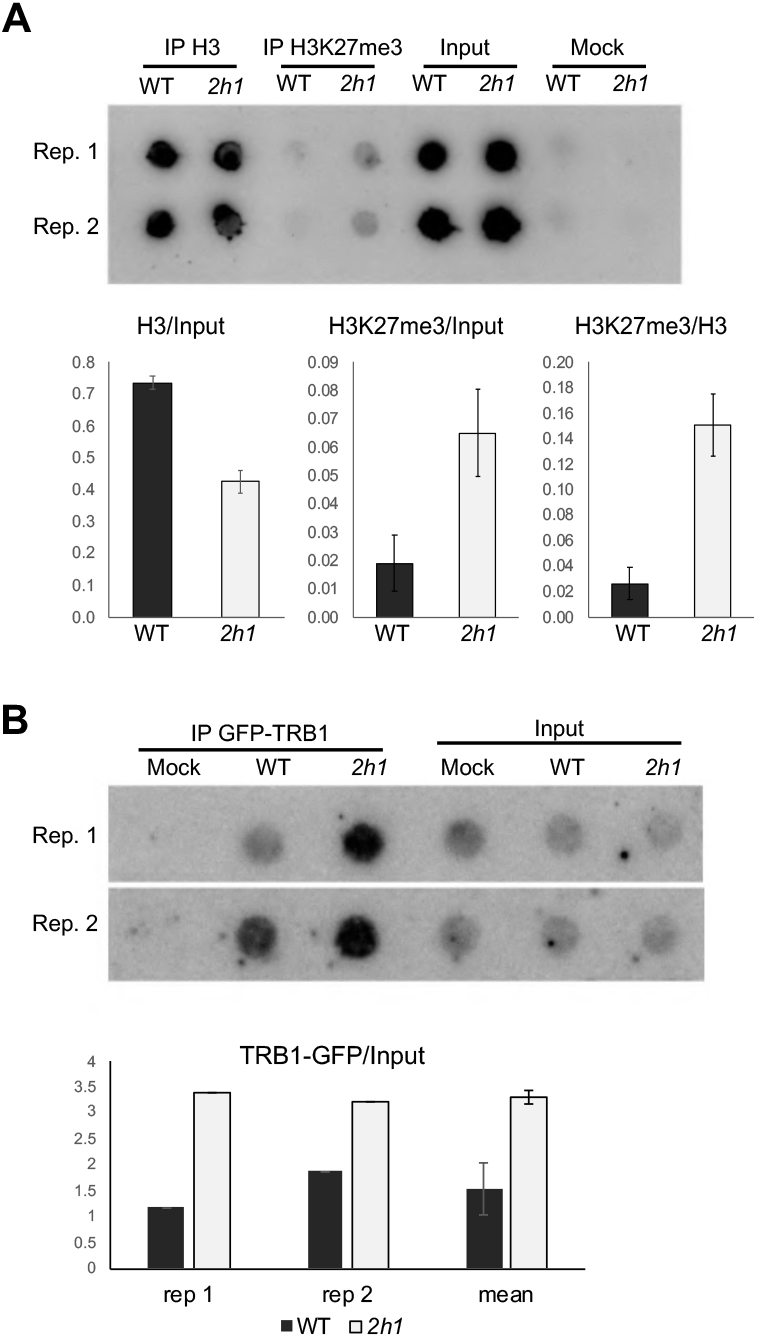
Figure 4 complementary data. **A.** Independent biological replicates of H3K27me3 and H3 ChIP DNA hybridization to telomeric probes completing Figure 4A. **B.** GFP-TRB1 in enriched at telomeres in *2h1* nuclei as compared to WT. Anti-GFP ChIP-blotting was performed as in Figure 4A and S4A using *TRB1::GFP-TRB1* and *2h1/TRB1::GFP-TRB1* plants. The results of two independent biological replicates are shown in the upper panel, and corresponding signal quantification in the lower panel.

**Figure S12:**
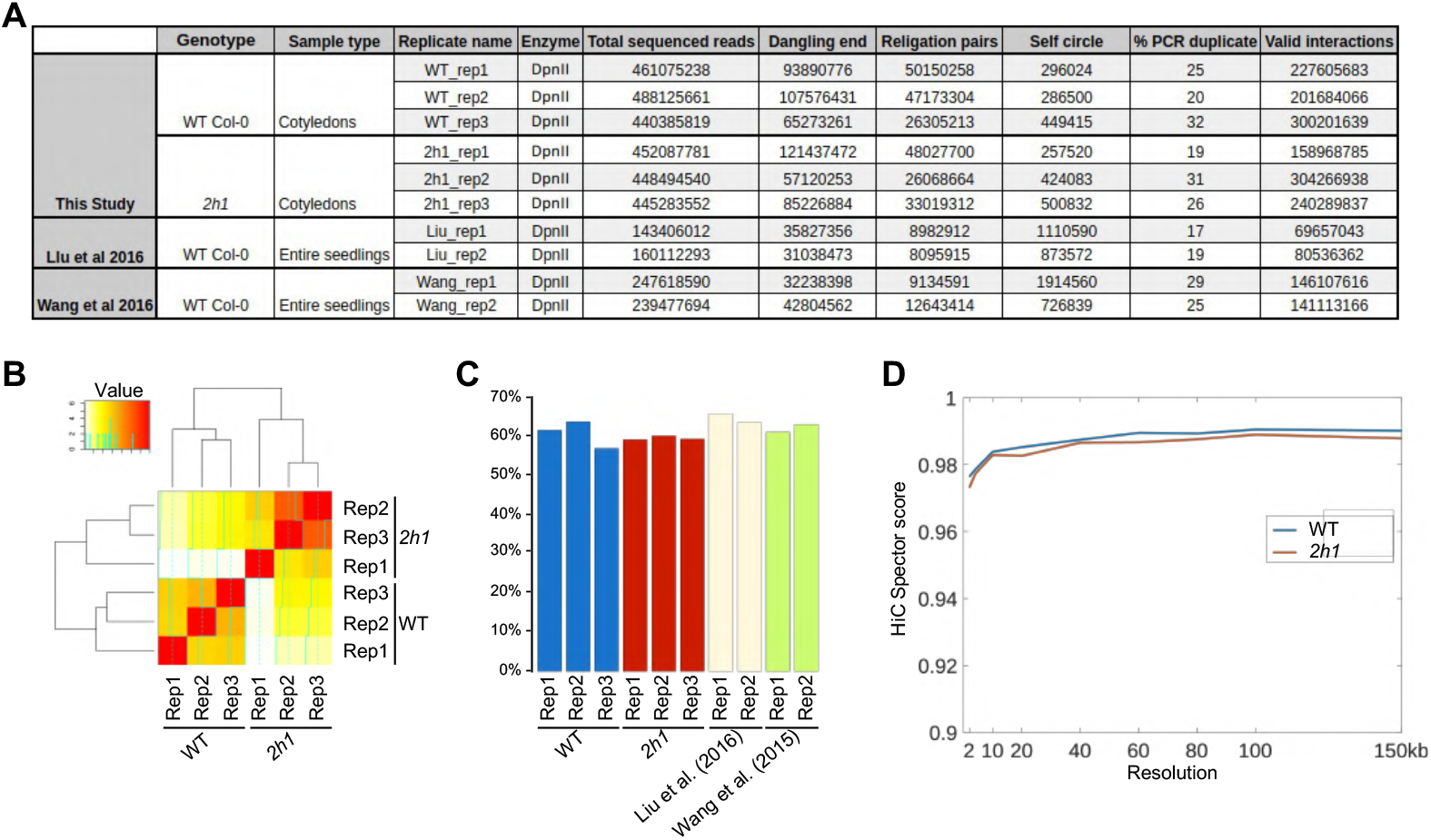
Hi-C mapping and similarity index quality. **A**. Mapping results of Hi-C libraries and comparison with re-processed Hi-C datasets from Liu et al. (2016) and Wang et al. (2015). **B**. Heatmap of similarity scores among WT and *2h1* mutant datasets calculated using Spearman rank correlation between each sample as 100 kb bins. **C**. Comparison of intra-chromosomal reads over the total number of valid interactions among our samples and published DpnII-based Hi-C datasets from Liu et al. (2016) and Wang et al. (2015). Proportion of *cis* (intra-chromosomal) interactions among valid interactions is positively correlated with library quality for *in situ* Hi-C as in Sun et al. (2020). **D.** Estimation of the Hi-C resolution achieved in this study. The curves show the Hi-C Spector score of chromosome 1 contact map down-sampled at 10% of contacts as in Carron et al. (2019). We computed 30 down-samples at a resolution of 2kb and computed Hi-C Spector score similarity, using an Eigen value of 10, against each Hi-C map with a resolution from 2kb to 150 kb. For each resolution, the maximum Hi-C Spector score of the 30 down-samplings is reported. This analysis was performed using the merge of three independent biological replicates. Public data sources are given in Additional file 3.

**Figure S13.**
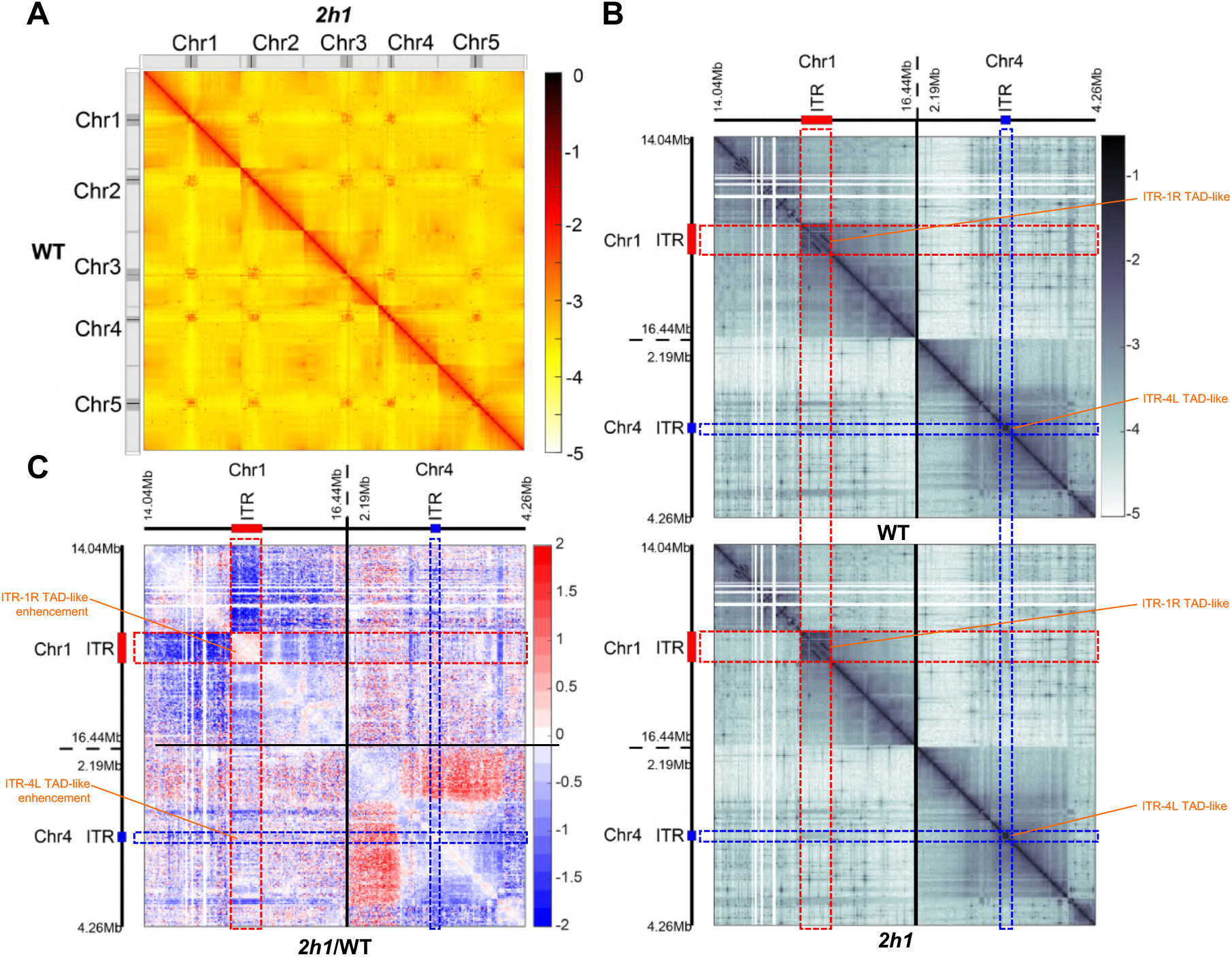
Chromosome and ITR-1R/4L topology in WT and *2h1* nuclei, completing Figure 5. **A.** Hi-C interaction frequency heatmap showing normalized Log_10_ contact count at 100 kb resolution for all *Arabidopsis* chromosomes in WT (below the diagonal) and *2h1* (above the diagonal) nuclei. Lateral tracks depict the positions of centromeres (black) and pericentromeres (gray). **B.** Same analysis than (A) illustrating interaction frequencies between the chromosomal regions spanning 1 Mb around ITR-1R (Chr1:15086191-15441067) and ITR-4L (Chr4:3192760-3265098) at a 10 kb resolution. **C.** Relative differences of interaction frequency between WT and *2h1* nuclei for the ITR regions illustrated in (B). Log_2_ ratios of normalized interaction frequencies of *2h1* vs WT nuclei are shown at a 10kb resolution. All Hi-C data were analyzed using the merge of three independent biological replicates.

**Figure S14.**
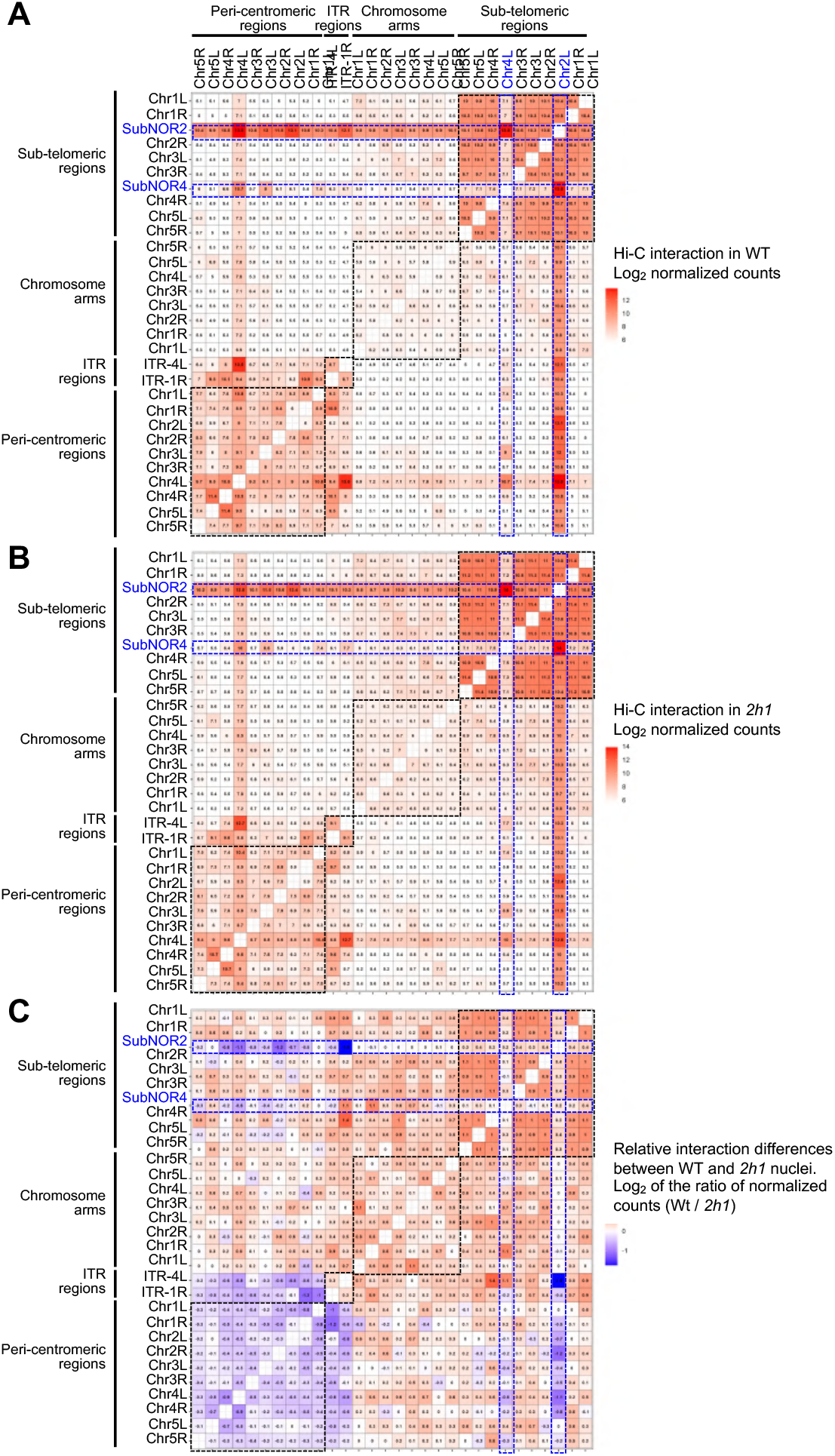
Frequency of interaction between pericentromeric, ITR-1R/4L, and telomeric regions in WT and *2h1* nuclei, and difference between the two genotypes. **A-B.** Pericentromere-embedded ITR-1R and 4L frequently associate with other pericentromeric regions through inter-chromosomal contacts, but less frequently with all other tested chromosome domains except the *NOR2* region. Besides frequent long-range interactions with *NOR4*, the *NOR2*-adjacent Chr2L region also shows frequent interactions with several genome regions including ITR-4L and its neighboring pericentromeric regions on Chr4L. **C.** Relative differences of interaction frequency. With the exception of *NOR*-proximal Chr2L and Chr4L regions, telomere-telomere and telomere-ITR interactions tend to increase in the absence of H1. Similarly, interactions between ITR-1R and 4L are slightly more frequent in the mutant line. In contrast, interactions between ITRs and all pericentromeres tend to be reduced in the mutant line. The decrease in frequency of interaction is particularly marked between the *NOR*-adjacent regions and the pericentromeres of chromosome 2 and 4. Interaction frequencies are expressed as logarithm of the observed read pairs (A and B) or logarithm of their ratio (C) normalized for region size. The indicated values are the median of three biological replicates (A and B) and the ratio of the medians (C). In (A-C) we probed four groups of regions: 1) ITR-1R and 4L coordinates, 2) sub-telomeric regions defined as the 100 kb terminal chromosomal regions adjacent to the telomeres, 3) pericentromeric regions represented by 100-kb segments located at 1 Mb from centromeres, and 4) 100-kb chromosome arm regions located at 5 Mb from the telomere positions. The sub-telomeric regions of chromosome 2 and 4 left arms are separated from the telomeres by the *NOR2* and *NOR4*, and therefore referred to as subNOR2 and SubNOR4 respectively (blue label). Distal arm regions were used as control.

**Figure S15.**
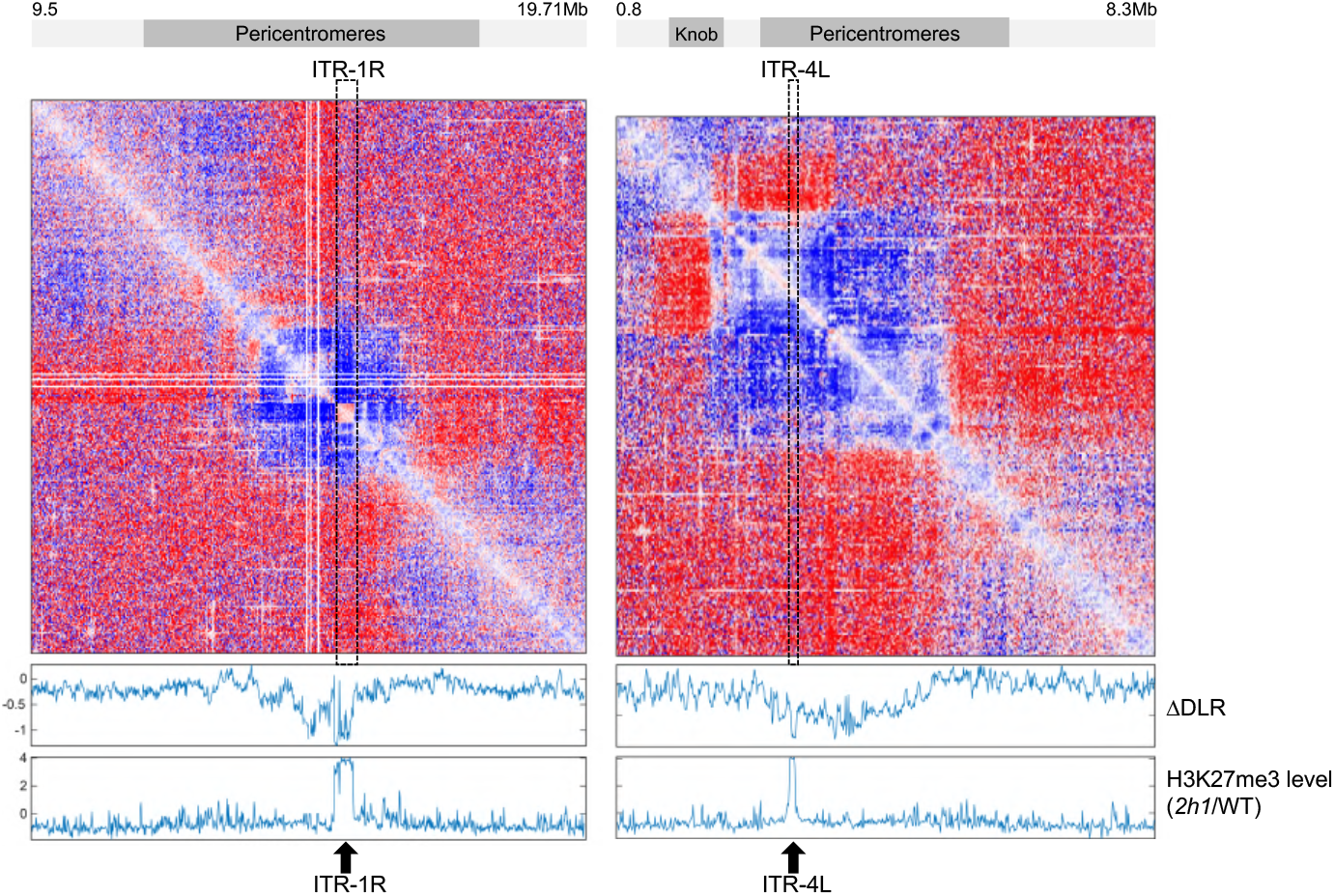
ITR1-1R and ITR-4L display lower DLR in *2h1* nuclei as compared to neighboring pericentromeric regions. The upper panel displays log2 ratio of O/E interaction frequency (*2h1*/WT) around ITR-1R and ITR-4L regions. Middle panel, ΔDLR represents the variations in distal-to-Local [log2] ratios in *2h1 vs* WT (*2h1*/WT) (see Methods). Bottom panel, ratio of H3K27me3 mean levels between *2h1* and WT nuclei (*2h1*/WT). Data combine three independent biological replicates and are processed at a 10kb resolution.

**Figure S16.**
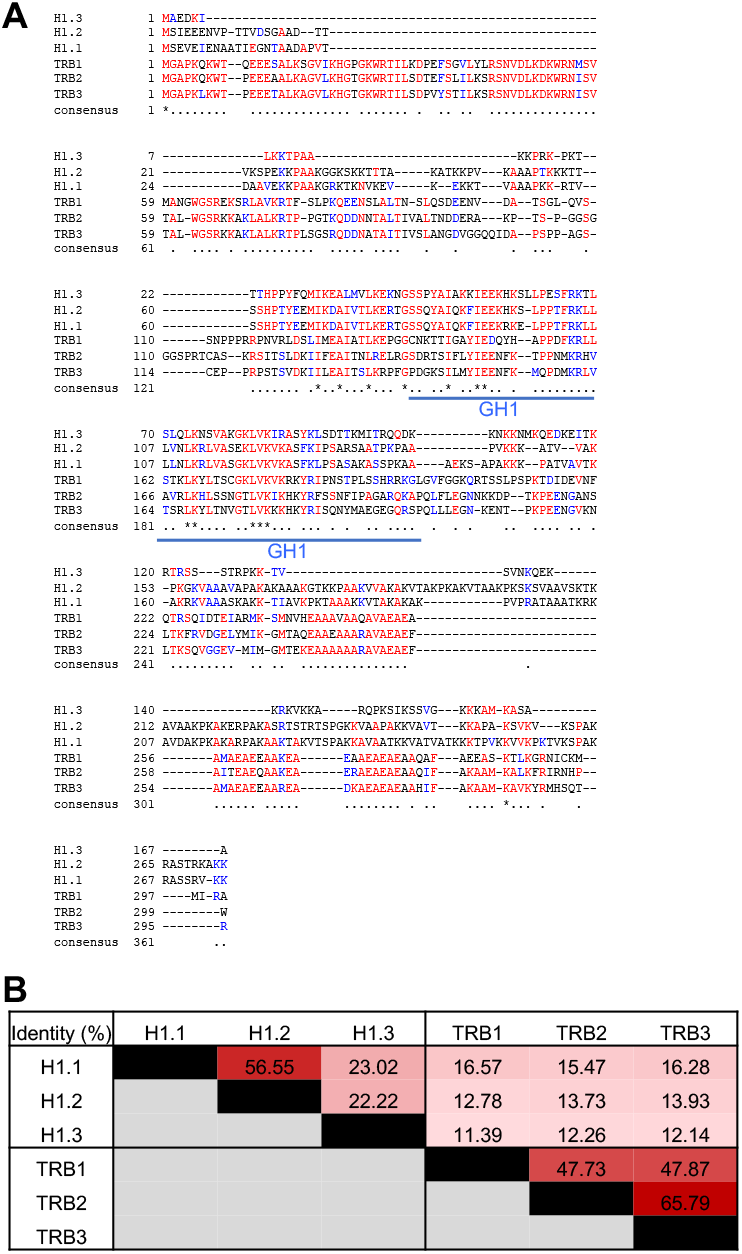
TRB family members display an amino-terminal Myb domain and a central GH1 domain that may explain their association to *telobox*-containing linker DNA. **A.** ClustalW protein sequence alignment of TRB1, TRB2 and TRB3 proteins with the three H1 variants showing the relative conservation of a central globular H1 domain. **B.** Amino-acids sequence identity between TRB1, TRB2 and TRB3 proteins and the three H1 variants.

**Figure S17.**
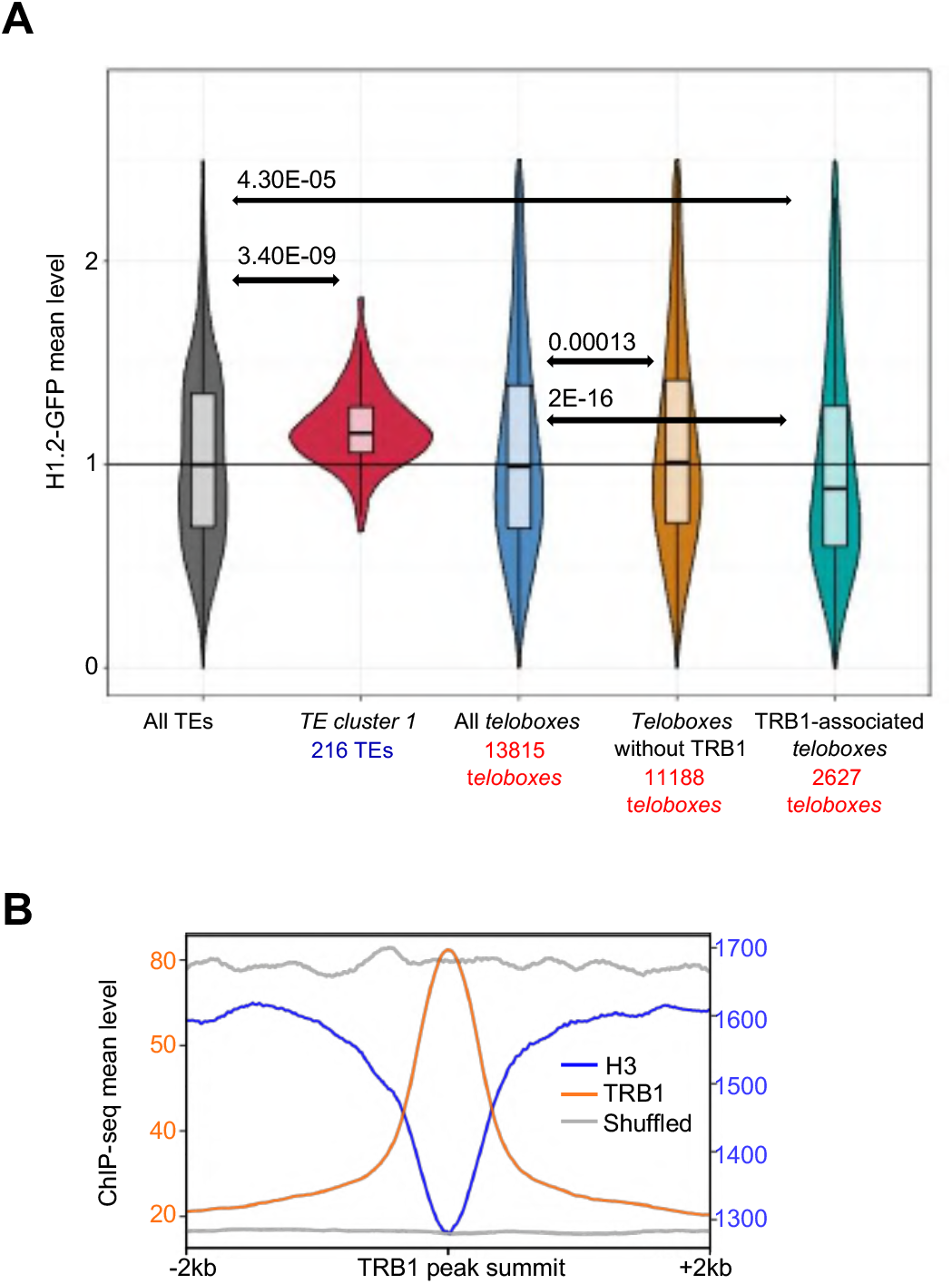
H1 and TRB1 display antagonistic chromatin association along the genome: complementary data to Figure 6. **A.** *TE Cluster 1* displays elevated H1 level as compared to the ensemble of *Arabidopsis* TEs and to other *telobox*-containing regions. In agreement with the proposed mutually exclusive binding of H1 and TRB1 over *teloboxes*, the set of *teloboxes* matching a known TRB1 peak displays significantly less H1 occupancy than other *teloboxes* of the genome. P-values of differences between the medians assessed using a Wilcoxon rank-sum test are shown. **B.** Figure 6D complementary data. H3 occupancy (read coverage) and randomly shuffled peaks are shown as control of data normalization. The shuffled control was produced with random permutations of genomic position of the regions of interest. TRB1 ChIP-seq data are from Schrumpfová et al. (2014), data sources given in Additional file 3.

**Figure S18.**
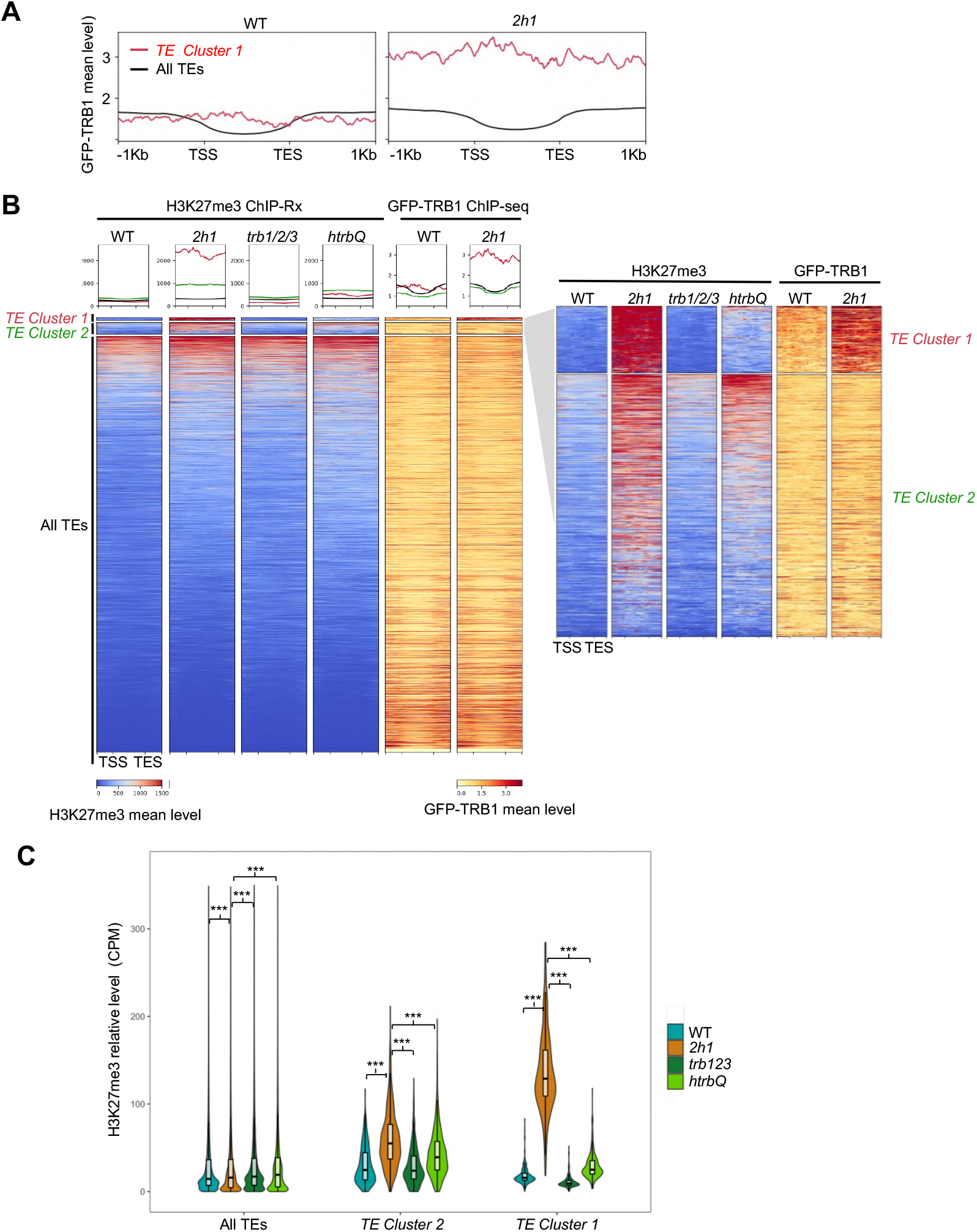
Figure 7 complementary analyses. **A.** GFP-TRB1 level at the indicated TE categories in WT and *2h1* nuclei (mean normalized coverage from two independent biological replicates). **B.** H3K27me3 and GFP-TRB1 mean level of different TE categories in the indicated genotypes. In each heatmap, TEs were ranked from top to bottom according to H3K27me3 or GFP-TRB1 mean level after RPCG or spike-in based normalization, respectively. While GFP-TRB1 ChIP-seq were performed using parental lines, H3K27me3 ChIP-Rx was performed on WT, *2h1* and *trb123* mutant lines selected from null F2 segregants from the same cross than the analyzed *2h1trb1trb2trb3* (*htrbQ*) plant line. **C.** Same analysis than (B) displaying H3K27me3 mean level over whole annotations TEs. All data represent the mean of two independent biological replicates.

